# A nematic framework for sprouting angiogenesis

**DOI:** 10.1101/2025.11.12.688071

**Authors:** Sara Barrasa-Ramos, Carles Blanch-Mercader, Abdul I. Barakat

## Abstract

Sprouting angiogenesis is critical for embryogenesis, wound healing, and tumor growth. Here we show that the liquid crystal framework, which has recently been leveraged to study various morphogenetic events, provides unique insight into angiogenic sprouting. In response to vascular endothelial growth factor, a potent pro-angiogenic factor, endothelial cells cultured on the surfaces of soft collagen hydrogels form elongated cell streams that coexist with more polygonal cell regions. Angiogenic sprouting initiates preferentially at cell stream tips, where nematic order gradients are most pronounced. We demonstrate that cell streams are associated with large traction forces that lead to 3D deformations of the underlying hydrogel and that stream formation results from a delicate balance between microtubule polymerization and actomyosin contractility. Finally, we show that active control of nematic order allows modulation of sprout formation, highlighting the key role of EC orientation gradients in angiogenic sprouting and paving the way for new therapeutic avenues.

**Teaser:** A nematic perspective reveals that sprouting angiogenesis correlates with elevated gradients in endothelial cell orientation.

## Introduction

Sprouting angiogenesis, the process of formation of new microvessels from pre-existing vasculature (1), is essential for embryonic development, wound healing, and tissue reperfusion after ischemia (2–5). Angiogenic sprouting also plays an important role in disease, with tumor development as a prominent example (6). Research in the past decades has provided great insight into the signaling pathways that govern sprout initiation and growth (7–11). More recently, the role of mechanical forces in angiogenic sprouting has become increasingly recognized; however, the literature on the role of biophysical factors in sprout formation contains conflicting information, and the underlying mechanisms remain largely unknown (12).

The initial stages of angiogenic sprouting involve a 2D-to-3D transition whereby a capillary branch emerges from the planar endothelial monolayer and grows into the underlying extracellular matrix. This process is reminiscent of other morphogenetic events that have recently found an explanation within the framework of active liquid crystals including epithelial cell extrusion (13), myoblast mound formation (14, 15), myoblast crisscross bilayering (16), and hydra morphogenesis (17, 18). The common denominator in all of these processes appears to be the role of topological defects - specific distortions in nematic alignment configurations - as starting points for the emerging morphological features. Although this remains an area of active investigation, the mechanical stresses generated by cell activity in the regions surrounding topological defects seem to be responsible for the observed phenomena in most of these cases.

Vascular endothelial growth factor (VEGF), a potent agonist that promotes sprouting angiogenesis (19), has been shown to induce drastic changes in endothelial cell (EC) morphology. More specifically, *in vitro* studies have demonstrated that VEGF elicits pronounced elongation in a subset of ECs (20), resulting in the formation of spindle-like cell streams with nematic alignment that are surrounded by patches of cells with less elongated shapes. This scenario of co-existence of nematic domains with more isotropic regions has, to our knowledge, no precedent in the morphogenesis literature.

In the present work, we are interested in determining how the interplay between EC activity and the distortions in nematic EC alignment induced by VEGF may influence angiogenic sprout initiation. To this end, we have developed a series of *in vitro* systems that involve culturing ECs to confluence on the surfaces of soft collagen hydrogels and subsequently monitoring angiogenic sprout development into the hydrogel in response to VEGF addition. Using these systems, we demonstrate that nascent sprouts co-localize with wedge regions, which are located at the tips of EC streams and are characterized by high gradients in nematic orientational order. Using a combination of analytical models and traction force microscopy (TFM) experiments, we demonstrate that these wedge regions are associated with particularly elevated traction forces. Interestingly, we observe that EC streams near wedges correlate with out-of-plane (i.e. perpendicular to the endothelial monolayer) micron-scale deformations in the underlying collagen substrate, which are in qualitative agreement with those predicted for a linear elastic material subjected to cell-generated traction forces. We also show that cell elongation within streams is determined by a competition between microtubule (MT) polymerization and actomyosin contractility, with impaired elongation resulting in the formation of vortex-like topological defects and altered substrate deformation patterns. We finally show that modulating endothelial nematic order through the application of either geometric constraints or mechanical cues to the endothelial monolayer can be exploited to effectively exert control over angiogenic sprout formation.

## Results

### VEGF induces both angiogenic sprouting and changes in endothelial morphology *in vitro*

As already mentioned, VEGF is not only a potent pro-angiogenic factor, but it is also known to induce dramatic changes in the morphology of ECs. To examine the two phenomena simultaneously and to study possible links between them, we cultured human umbilical vein endothelial cells (HUVECs) to confluence on top of a flat, fibronectin-coated type I collagen hydrogel substrate with a thickness of 250 µm (figs. S1 and **1**A). We then quantified the effect of VEGF (50 ng*/*mL applied for 66 h) on both angiogenic sprouting and EC morphology.

Sprout identification was initially conducted under confocal microscopy. As such, cord-like multi-cellular structures extending into the collagen hydrogel substrate without detaching from the endothelial monolayer at their base were considered as sprouts. Sprouts were sometimes lumenized, and they were in some cases observed to also connect to the monolayer at their distal tip (fig. **1**B and S2). Manual quantification of sprout density was then performed on widefield image stacks, revealing that while sprouts were largely absent in control experiments, addition of VEGF significantly increased sprout density. For the VEGF concentration and incubation time considered, the sprout density in VEGF-treated ECs was determined to be 2.0 ± 1.2 sprouts*/*mm^2^ (mean ± SD, *n* = 4, fig. **1**C).

**Fig. 1.**
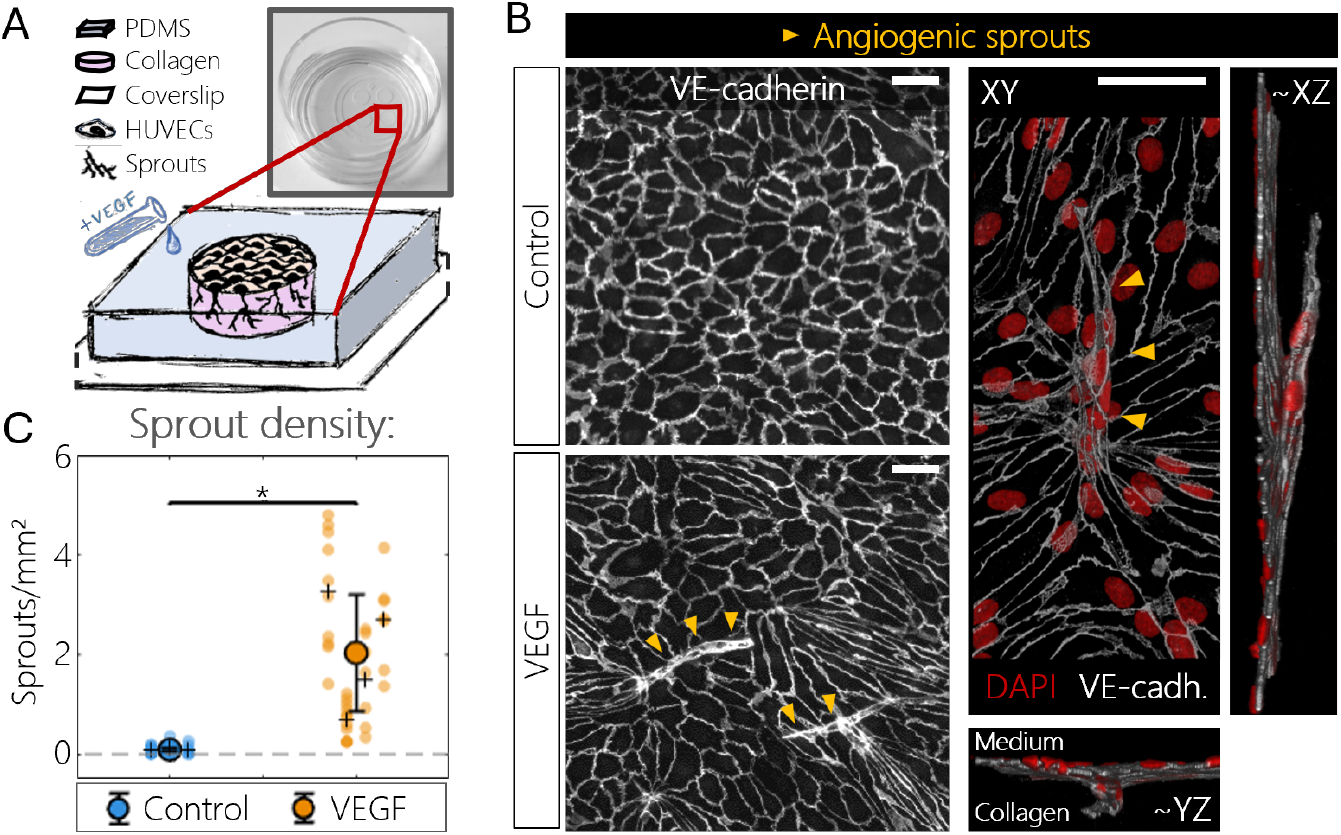
Effect of VEGF on angiogenic sprouting *in vitro*. **A:** Schematic of the employed experimental platform (not to scale). **B, left:** VE-cadherin (cell-cell junctions, white) immunofluorescence images reveal the formation of sprouts, demarcated by gold arrowheads, under VEGF conditioning and their absence in control conditions. **B, right:** Confocal reconstruction of a sprout (gold arrowheads) from immunostaining for VE-cadherin and DAPI (nuclei, red). Scale bars, 50 µm. **C:** Effect of VEGF (50 ng*/*mL for 66 h) on sprout density. Colored dots represent 8-12 replicates per experiment; black crosses represent the average per experiment. n=3–4 independent experiments, two-sample t-test (^∗^*P <* 0.05). Error bars represent ±SD.

In addition to triggering sprouting and consistent with previous reports (20, 21), VEGF induced morphological changes in the EC population. To quantify these changes, the cell inverse aspect ratio (minor-to-major axis ratio) was assessed (fig. **2**). While control ECs exhibited a broadly polygonal morphology with an average inverse aspect ratio of 0.63 ± 0.02 (mean ± SD, *n* = 3, fig. **2**B), addition of VEGF triggered significant elongation in a subset of the cells, skewing the distribution and lowering the mean inverse aspect ratio to 0.40 ± 0.01 (mean ± SD, *n* = 3, fig. **2**B). The spatial heterogeneity in EC morphology in VEGF-treated cells translated into the formation of cell streams, fascicles of highly elongated and aligned cells (cyan clusters in fig. **2**A) surrounded by regions of less elongated cells that resemble those under control conditions (purple-to-magenta regions in fig. **2**A). Although these less elongated cells outside the streams exhibit a certain degree of elongation, we will henceforth refer to them as ‘polygonal’ in order to contrast them with the more elongated cells within the streams.

**Fig. 2.**
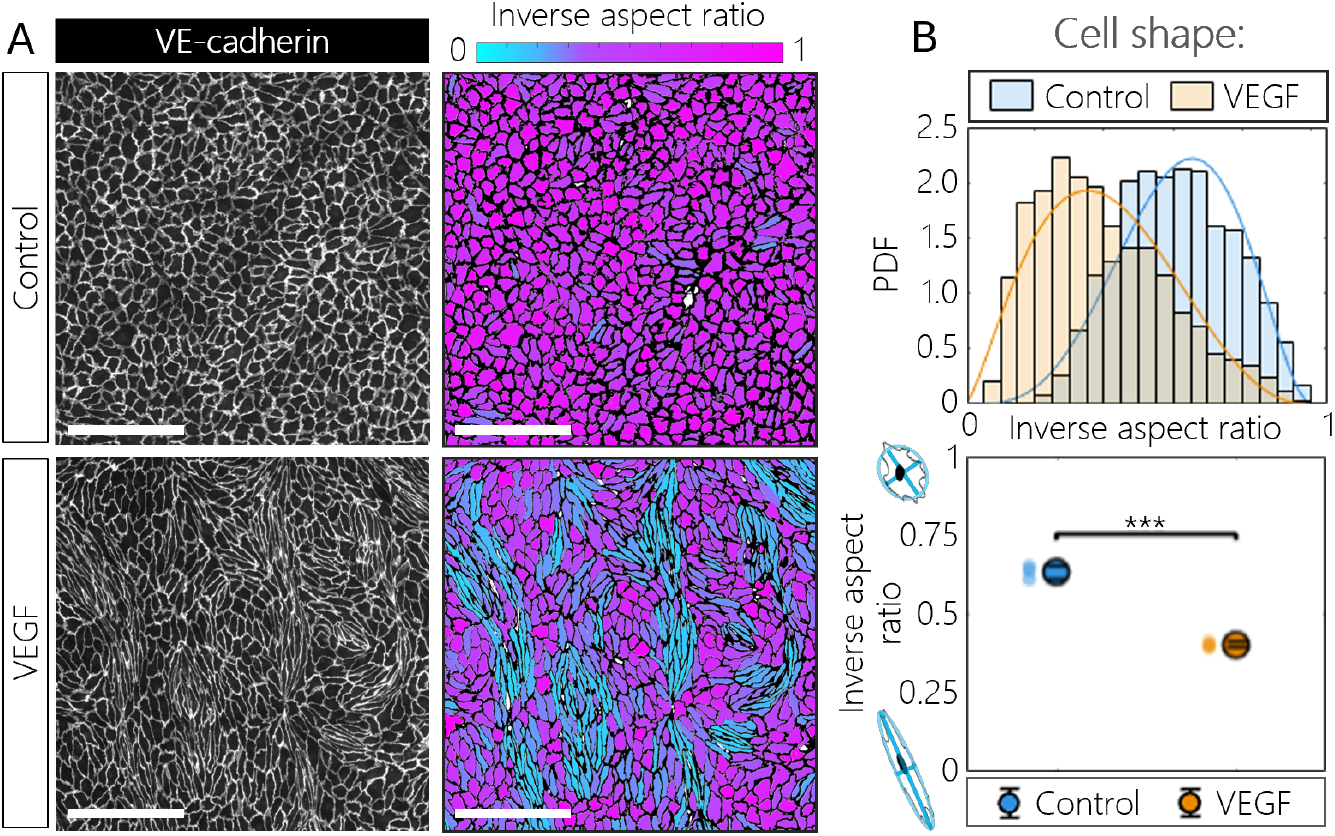
Effect of VEGF on endothelial cell shape *in vitro*. **A, left:** Widefield images from immunostaining for VE-cadherin (cell-cell junctions, white) of either control or VEGF-conditioned monolayers. **A, right:** Color coded inverse aspect ratio (minor-to-major axis ratio) of the cells in the leftmost images. Scale bars, 250 µm. **B, top:** Representative distributions of cell inverse aspect ratio are fitted by a beta distribution of the first kind (solid curves). **B, bottom:** Quantification of the average inverse aspect ratio per monolayer shows increased elongation under VEGF conditioning (50 ng*/*mL for 66 h). Data from n=3 independent experiments, one-way ANOVA, Dunnett’s post-test (^∗∗∗^*P <* 0.001). Error bars represent ±SD.

### Nascent sprouts co-localize with wedge regions

Sprouts were observed to typically emerge along the boundaries of cell streams (figs. S3 and **3**A). To explore a potential link between the spatial heterogeneity in EC morphology and sprouting angiogenesis, we analyzed the sprout locations relative to the cell streams. We were able to identify specific zones along stream boundaries where cells within the stream aligned perpendicular to the boundary, and we termed these zones “wedges” (see Materials and methods and fig. **3**A). Distance maps to wedges could then be calculated for each region and for sprouts (fig. **3**A). Comparison of the distribution of sprout-to-wedge distances (white distribution, fig. **3**B) with the distributions of distances between wedges and all points within the cell stream and polygonal cell regions (cyan and pink distributions, fig. **3**B) revealed a degree of co-localization between sprouts and wedges. In spite of the area imbalance, this result is independent of the type of region considered (cell streams or polygonal cell regions, fig. **3**B), showing that wedges in particular, and not cell streams more generally, are preferential sites for sprouting angiogenesis.

**Fig. 3.**
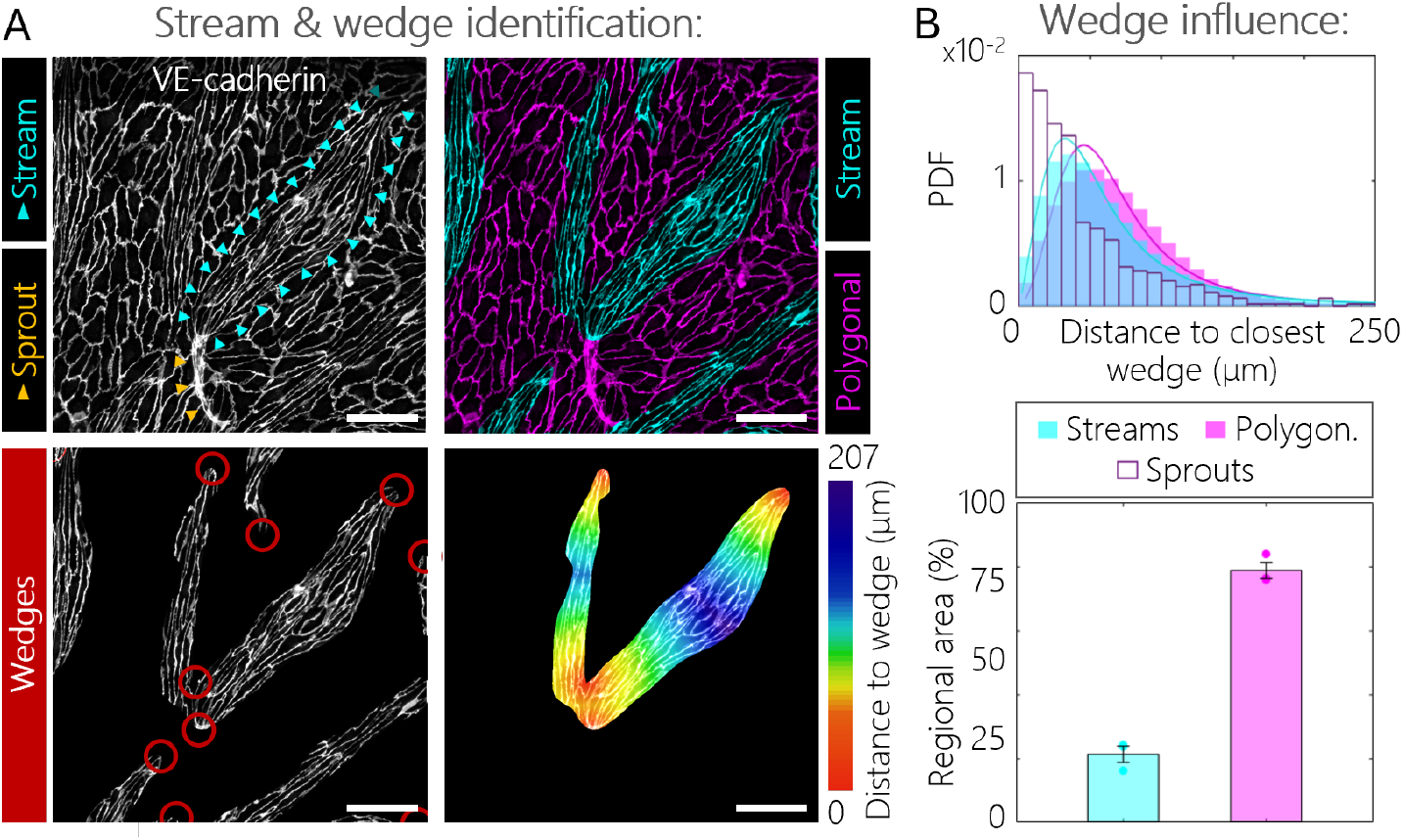
Wedges are preferential sites for sprouting angiogenesis. **A, left-top:** Immunostaining for VE-cadherin (cell-cell junctions, white) showing a sprout (yellow arrowheads) associated with a cell stream (cyan arrowheads) under VEGF conditioning. **A, right-top:** Segmentation of cell stream (cyan) and polygonal cell (magenta) regions. **A, left-bottom:** Wedges, within red circles, as defined by eq. 2 in Materials and methods. **A, right-bottom:** Example of distance map to nearest wedge in a specific region. Scale bars, 100 µm. **B, top:** Probability distributions for the distance to the regional closest wedge for sprouts (white, n=647) as well as for cell stream (cyan) and polygonal cell (magenta) regions. Pooled data from n=4 independent experiments. **B, bottom:** Percentage of area occupied by each region type (stream or polygonal). Data from n=3 independent experiments.

### Wedges represent singular mechanical environments

The results above pointed to a link between distortions in nematic cell alignment in the EC monolayer and angiogenic sprouting. Previous theoretical studies in active nematic models have shown that specific stress patterns arise in regions surrounding topological defects, which are singular configurations of the nematic field (22, 23). Furthermore, experimental studies have shown that angiogenic sprouts can deform the underlying matrix as they grow (24). These observations led us to hypothesize that the VEGF-induced alterations in EC morphology might be accompanied by specific traction force patterns leading to deformations of the collagen substrate.

To investigate this possibility, we first analyzed the crystalline order of both control and VEGF-conditioned ECs and found that in both cases, the nematic order was predominant at all scales up to 200 µm (fig. S4A). Active nematics can generate traction force densities **t** via an interplay between distortions of the nematic director field **n** and activity. Based on symmetry arguments, three possible force densities with nematic symmetry are expected in two-dimensional geometries (25, 15, 26), which take the form:

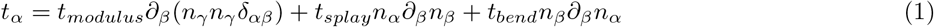

To account for the variations in EC shape, the norm of |**n**| was allowed to vary in space (see Materials and methods). Each force term in eq. 1 depends on the director field and its gradients up to a constant material parameter *t*_*modulus*_, *t*_*splay*_ or *t*_*bend*_ that is proportional to the cellular activity level. While the term proportional to *t*_*modulus*_ represents traction forces that arise from spatial variations in EC shape, the terms proportional to *t*_*splay*_ and *t*_*bend*_ capture traction forces that result from gradients in cell alignment. More specifically, splay encodes the divergence of the director field, while bend is associated with its curvature, as illustrated in fig. S5. By determining the director field on EC monolayers **n** (see Materials and methods), we computed each of these forces in ECs. While the traction force patterns were mostly uniform under control conditions, their magnitude peaked near the wedges in VEGF-treated monolayers, thereby predicting increased traction forces at these locations (fig. S6).

To test the prediction of traction force distribution described above, we performed TFM experiments under VEGF treatment by introducing fluorescent polystyrene beads into the underlying collagen substrate (see Materials and methods). The resulting averaged traction force maps (fig. **4**B) displayed a magnitude peak in the wedge region and forces that were primarily directed towards the cell stream bulk. These features could be reproduced by eq. 1 (fig. **4**C). Furthermore, a distribution mimicking the traction force map in fig. **4**B was able to reproduce the in-plane displacement field on the boundary of a linear elastic semi-infinite region representing the collagen hydrogel (figs. **4**B and **4**C).

**Fig. 4.**
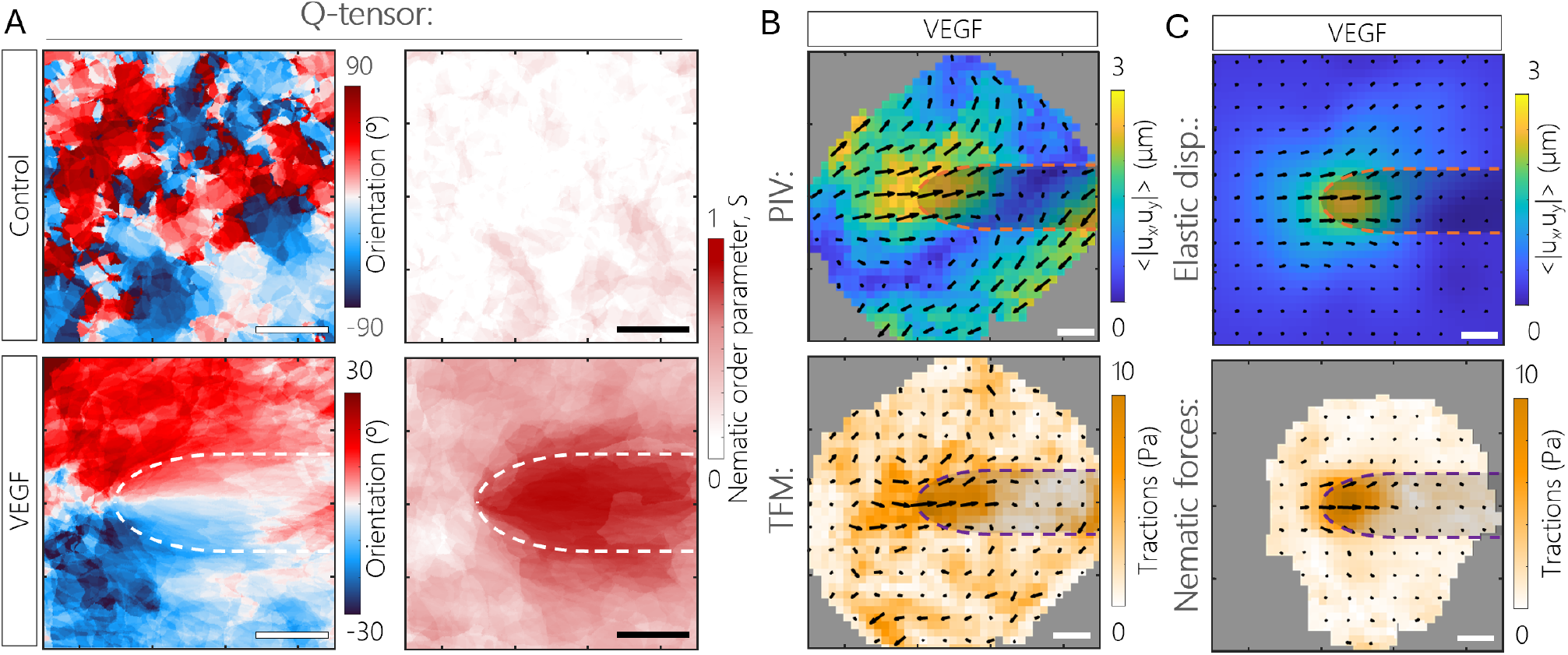
Nematic model predicts traction forces at the wedge that are compatible with 2D TFM experiments. **A:** Average orientation angle (left panels) and nematic order parameter (right panels) of control experiments and around wedges in VEGF-conditioned experiments (see Materials and methods). Pooled data (n_control_=13 and n_VEGF_=17) from n_control_=4 and n_VEGF_=7 independent experiments. **B, top:** 2D PIV map shows average hydrogel displacements around wedges. **B, bottom:** Average TFM results reveal the traction force map around wedges. Pooled data (n=17 wedges) from n=3 independent experiments. **C, bottom:** The TFM force peak around the wedge can be captured by eq. 1. The computation of the nematic traction force map using eq. 1 is explained in Materials and methods. **C, top:** In-plane displacement field on the boundary of a half-space linear elastic material resulting from the application of Gaussian loads mimicking the traction force map in **C, bottom** (see Materials and methods). Scale bars, 50 µm. In all plots, the arrows show direction and relative magnitude, and the background is color-coded by absolute magnitude. Dashed lines and shaded regions indicate the approximate location of cell streams.

### VEGF conditioning induces substrate wrinkling

Next, we hypothesized that the mechanical forces themselves may be directly involved in the out-of-plane transition that occurs at the onset of sprout formation. To explore this possibility, we began by using a model that describes the collagen substrate as a linear elastic semi-infinite region subjected to Gaussian loads on its upper boundary (see Materials and methods). Considering a literature-based Poisson’s ratio for collagen hydrogels of 0.3 (27–29), this model closely reproduced the experimentally observed in-plane displacement map when loads were distributed so as to match the traction force density mimicking the TFM results (figs. **4**B and **4**C, see Materials and methods). Notably, the model also predicted an out-of-plane displacement of the collagen surface in the wedge region (fig. **5**A).

**Fig. 5.**
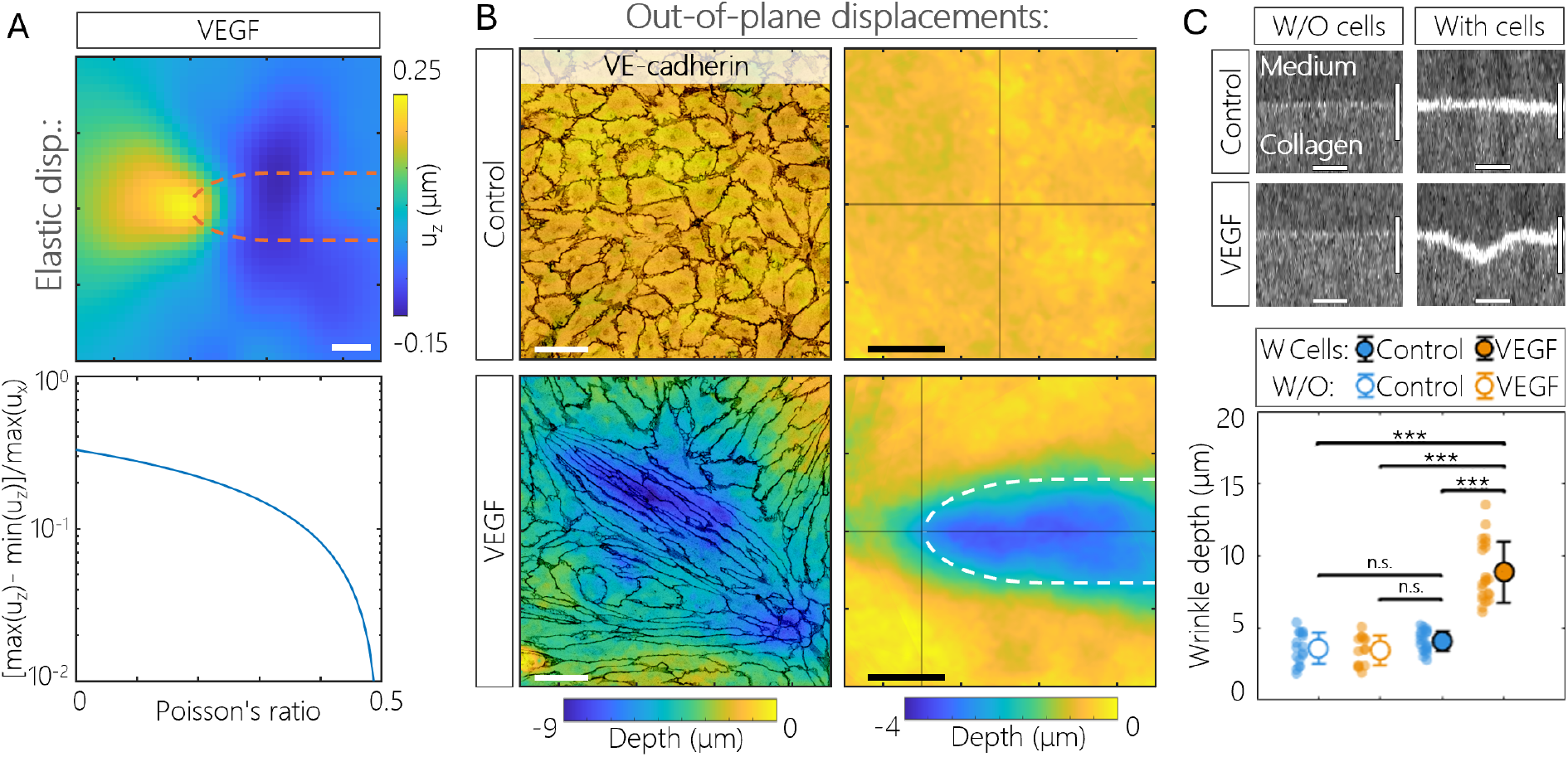
Substrate wrinkling occurs at the level of cell streams. **A, top:** Out-of-plane displacement field on the boundary of a half-space linear elastic material resulting from the application of Gaussian loads mimicking the traction force map in fig. **4**C (see Materials and methods). **A, bottom:** For Poisson’s ratios between 0 and 0.5, the linear elastic model predicts in-plane displacements *u*_*x*_ to be larger than out-of-plane displacements *u*_*z*_ derived from the traction force map in fig. **4**C (see Materials and methods). **B, left:** Example height maps of endothelial monolayers under control and VEGF conditioning. Cell contours are shown through VE-cadherin immunostaining (black). **B, right:** Average height maps reveal consistent wrinkle formation under VEGF-induced cell streams. The white dashed line indicates the approximate location of cell streams. No wrinkles are observed in the absence of VEGF. Scale bars, 50 µm. **C, top:** OCT images of the collagen surface reveal that both cell presence and VEGF conditioning are necessary for wrinkle formation. The cells appear as a white layer on the top surface of the collagen hydrogel. **C, bottom:** Wrinkle depth under control and VEGF treatment both in the presence and absence of cells. Pooled data from n=3 independent experiments, Kruskal-Wallis, Dunn’s post-test (^∗∗∗^*P <* 0.001, ^*n*.*s*.^*P >* 0.05 represents statistical non-significance). Error bars represent ± SD. Horizontal scale bars, 50 µm; vertical scale bars, 20 µm.

To test this prediction, we imaged the surface of the EC-covered collagen substrate using confocal microscopy. These observations revealed the presence of wrinkles in the collagen surface (valleys in the landscape) in VEGF-treated ECs (fig. **5**B). Consistent with our model, these wrinkles localized preferentially under cell streams. Quantification of wrinkle depth using optical coherence tomography (OCT) confirmed an increase in wrinkle depth in VEGF-conditioned samples (8.9 ± 2.1 µm; *n* = 22) relative to control samples (4.1 ± 0.7 µm; *n* = 21) (fig. **5**C), where substrates were largely flat. Nevertheless, two aspects of the experimentally observed out-of-plane displacements differ from our theoretical results. First, the substrate indentation occurs along the entire length of the cell stream rather than being localized near the wedge (figs. **5**A and **5**B). Second, for materials with a Poisson’s ratio between 0 and 0.5, the magnitude of the out-of-plane displacements predicted by the model are smaller than the in-plane displacements (figs. **4**C and **5**A); however, we experimentally observe them to be of the same order of magnitude (figs. **4**B and **5**B). While the first discrepancy may be explained by potential deviation of the collagen material properties from the linear elastic behavior assumed in the model including the possible effects of plasticity (30–32), the second discrepancy suggests the potential involvement of other mechanisms in wrinkle formation.

### Differential matrix degradation, but not deposition, contributes to wrinkle formation

As a first step in exploring alternative wrinkle formation mechanisms, we confirmed by OCT imaging that substrate wrinkling required the presence of an EC monolayer as evidenced by the fact that the surfaces of bare collagen substrates remained flat with and without VEGF (fig. **5**C). Therefore, VEGF in itself does not induce surface wrinkling of the collagen hydrogel. Subsequently, we hypothesized that the wrinkling of the collagen surface could be driven by either a local increase of basement membrane (BM) protein deposition outside the cell streams and/or a localized increase of extracellular matrix degradation within the cell streams. To test the protein deposition hypothesis, we stained fixed monolayers for the three BM proteins laminin, collagen IV, and fibronectin and measured the BM protein thickness (see Materials and methods). No differences in BM thickness were observed between the wrinkled and the non-wrinkled regions, suggesting that differential matrix deposition does not contribute to wrinkle formation (figs. **6**A and S7). To test the differential matrix degradation hypothesis, two types of analysis were carried out. First, fixed monolayers were stained for membrane type 1-matrix metalloproteinase (MT1-MMP), and staining intensity was compared between wrinkled and non-wrinkled regions. MT1-MMP density was higher in the cell stream regions, suggesting a role for this membrane-bound metalloproteinase in the formation of wrinkles (fig. **6**B). For the second analysis, alongside VEGF conditioning, broad spectrum MMP inhibition via the application of Ilomastat had no effect on wrinkle depth (fig. **6**C). This result, together with the small percent difference in MT1-MMP expression between the wrinkled and non-wrinkled regions (*<*5 %), suggests that differential matrix degradation is a minor contributor to wrinkle formation.

**Fig. 6.**
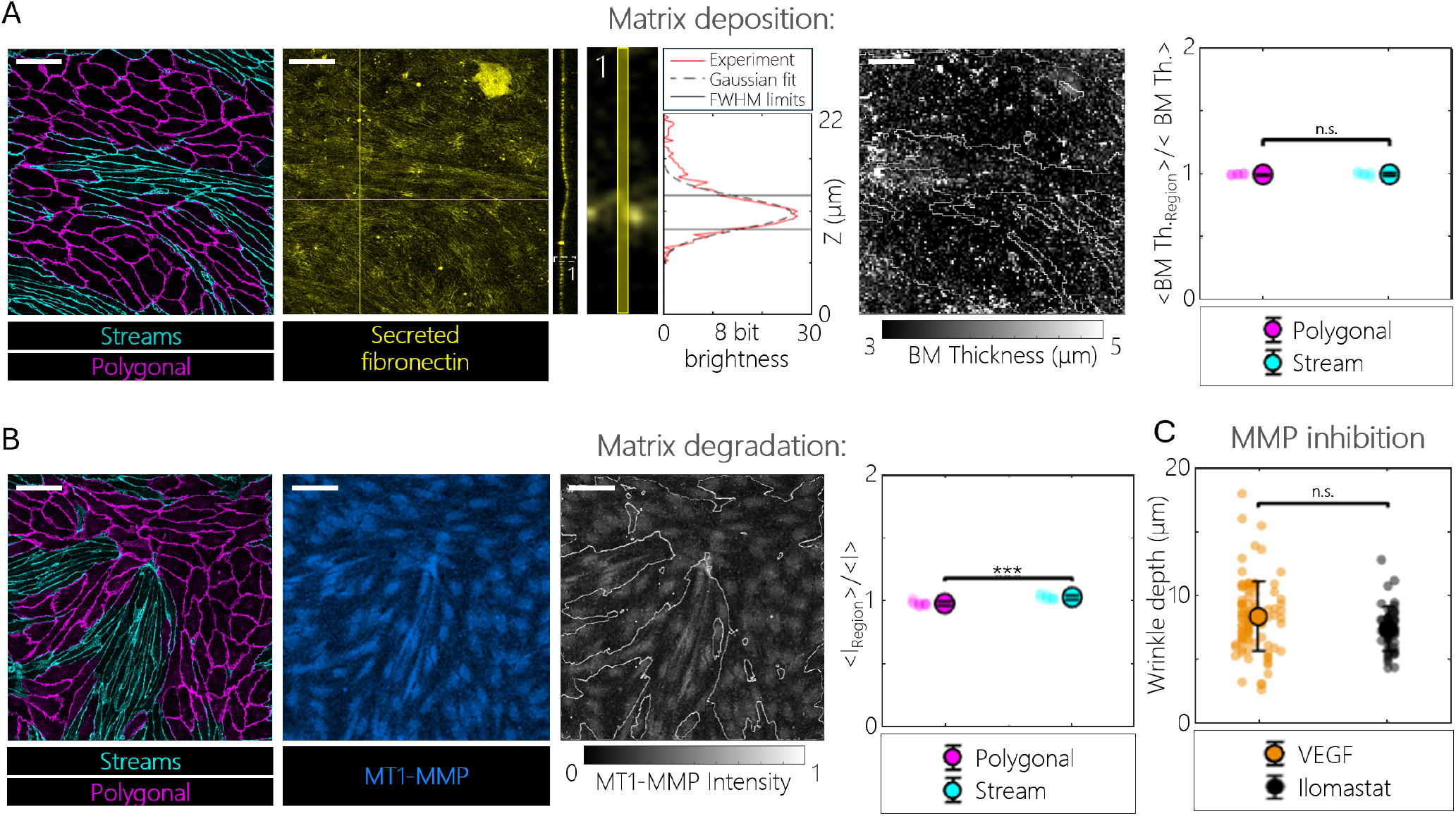
Role of ECM deposition and degradation in wrinkle formation. **A, from left to right:** stream-polygonal region segmentation. Corresponding immunostaining for secreted fibronectin (basement membrane component, yellow) and cross-section (solid yellow lines indicate the cross-sectional planes). Secreted fibronectin thickness map obtained from confocal stacks by calculating the full width at half maximum (FWHM) of a Gaussian fit of the Z-wise intensity for every XY coordinate. Thickness calculation example from Zone 1 in fibronectin immunostaining. Quantification of the average regional thickness normalized by the total image average shows no significant difference among regions. Pooled data (n=9) from n=3 independent experiments, two-sided Wilcoxon rank sum test (^*n*.*s*.^*P >* 0.05 represents statistical non-significance). **B, from left to right:** stream-polygonal region segmentation. Corresponding immunostaining for MT1-MMP (blue). Corresponding 8-bit maximum intensity projection of the MT1-MMP confocal stack normalized by 2 × 10^8^. Quantification of the average regional intensity normalized by the total image average indicates higher MMP concentration at the cell stream level. Pooled data (n=11) from n=3 independent experiments, two-sided Wilcoxon rank sum test (^∗∗∗^*P <* 0.001). **C:** Broad spectrum inhibition of MMPs does not significantly alter wrinkle depth. Pooled data from n=3-7 independent experiments, Kruskal-Wallis, Dunn’s post-test (^*n*.*s*.^*P >* 0.05 represents statistical non-significance). Errorbars represent ± SD. Scale bars, 50 µm.

### The role of the cytoskeleton in cell stream formation

Previous studies have suggested that cell shape is governed by a balance between the dynamics of microtubule (MT) growth and actomyosin contractility (33–35). Based on this notion, we anticipated that actin filaments and MTs would exhibit increased alignment within the cell stream regions of VEGF-treated cells. Quantifying the orientation of these cytoskeletal networks revealed that both actin and MT structures remain largely isotropic under control conditions, with MTs primarily organized radially around a central microtubule-organizing center (MTOC) (zone 1, fig. **7**A), while actin formed aster-like stress fibers connecting adjacent cells (zone 1, fig. **7**B). VEGF treatment induced alignment of both MTs and actin within the cell streams. More specifically, MTs organized into two diametrically opposed bundles around the MTOC (zone 3, fig. **7**A), and long actin stress fibers were accompanied by an increase in cortical actin (zone 3, fig. **7**B). Conversely, in the polygonal cell regions, both MTs and actin stress fibers exhibited a more spread distribution, recovering an organization similar to that observed under control conditions (zones 2, figs. **7**A and **7**B). These results demonstrate that cytoskeletal organization within cell streams is dramatically different from that within polygonal cell regions.

**Fig. 7.**
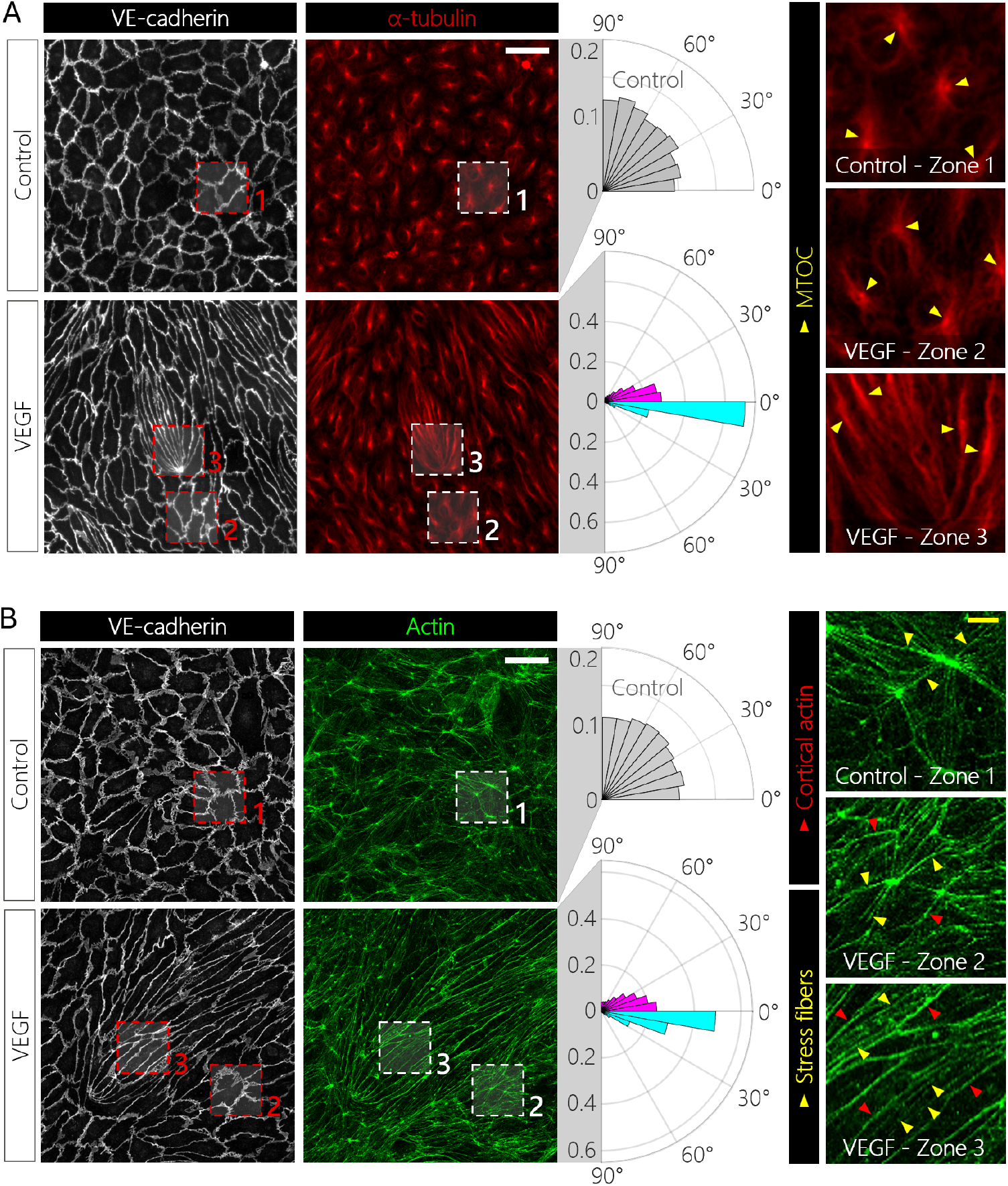
Organization of the cytoskeleton in cell streams vs. polygonal cell regions. **A: Microtubules** – VE-cadherin (white) and *α*-tubulin (microtubules, red) immunostaining of control and VEGF-conditioned monolayers. Histograms show the orientation angle distributions (absolute value) of *α*-tubulin in control monolayers (grey) and in cell stream (cyan) and polygonal cell (magenta) regions of VEGF-conditioned monolayers relative to the stream axis. Details of the three zones are shown on the right. Yellow arrows point to the MTOC of different cells. **B: Actin** – VE-cadherin (white) and phalloidin (F-actin, green) immunostaining of control and VEGF-conditioned monolayers. Histograms show the orientation angle distributions (absolute value) of control monolayers (grey) as well as cell stream (cyan) and polygonal cell (magenta) regions of VEGF-conditioned monolayers relative to the stream axis. Details of the three zones are shown on the right. Arrowheads point to cortical actin (red) or stress fibers (yellow). Pooled data (n=6-19 regions) from n=2-7 independent experiments. White scale bars, 50 µm; yellow scale bars, 10 µm.

### Cell stream formation and wrinkle depth are governed by a balance between MT dynamics and actomyosin contractility

Based on our observations of the organization of MT and actin networks and in light of previous studies implicating the EC cytoskeleton in angiogenic activity (36, 37), we investigated the impact of various cytoskeleton-altering agents on the formation of both cell streams and substrate wrinkles. In line with previous findings (20), MT dynamics proved to be key for cell elongation induced by VEGF treatment, with both MT depolymerizing (nocodazole) and stabilizing (taxol) agents resulting in a striking loss of cell elongation (fig. **8**A.a). Increasing actomyosin contractility with calyculin A had a similar effect on cell shape, leading to more homogeneous cuboidal monolayers, while decreasing actomyosin contractility with blebbistatin had no effect on cell elongation (fig. **8**B.a). Increased cell elongation was also observed upon disruption of actin stress fibers with cytochalasin D and latrunculin A (fig. **8**B.a). These results suggest that a specific balance between the actomyosin machinery and MT dynamics is necessary for the co-existence of cell streams and polygonal cell regions in VEGF-treated cells, with MTs promoting cell elongation and the actomyosin machinery counteracting this effect.

**Fig. 8.**
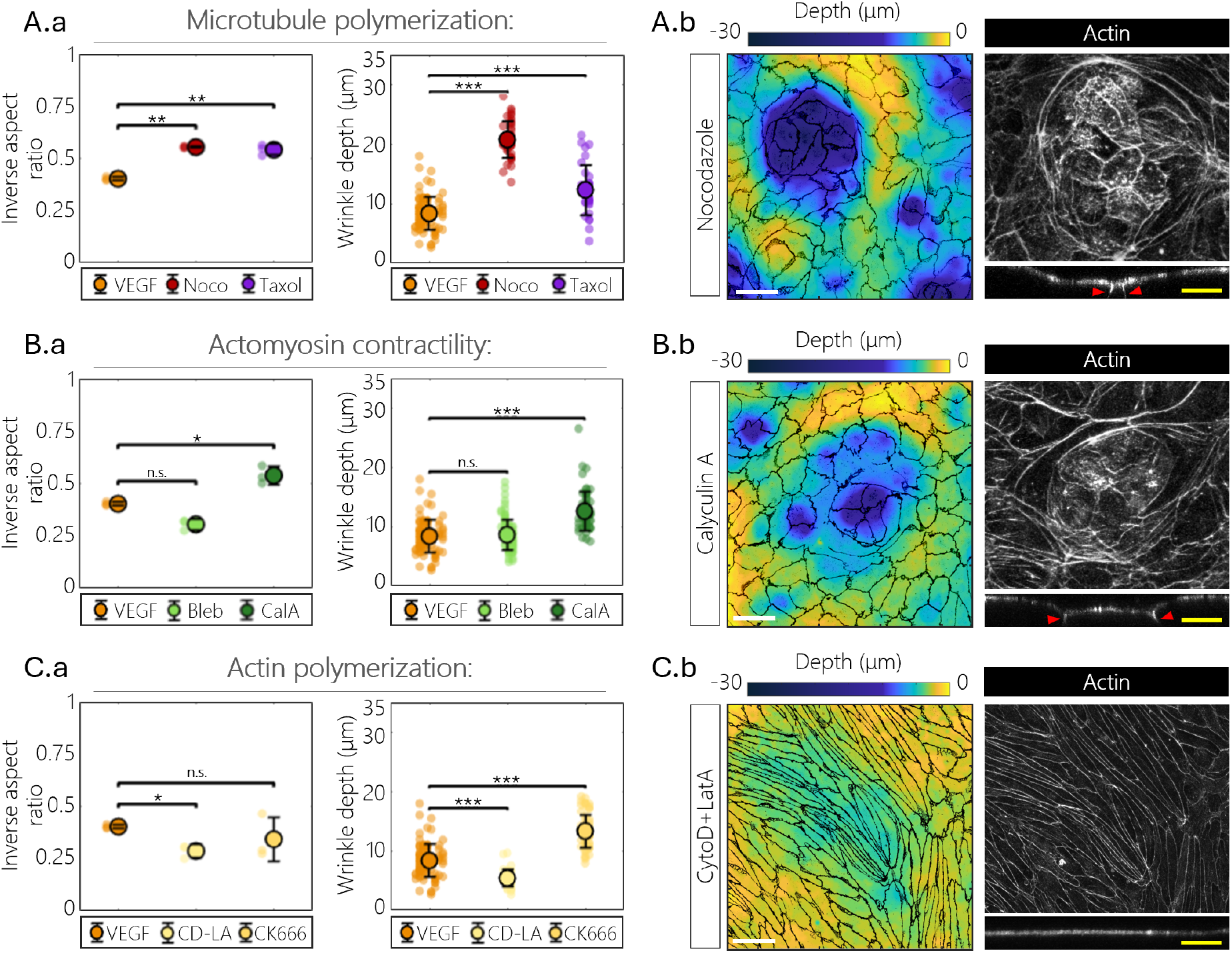
Role of cytoskeletal elements in cell stream and wrinkle formation. From left to right in each panel: quantification of inverse aspect ratio and wrinkle depth, examples of height maps, and immunostaining for actin (phalloidin, grey) of monolayers treated for 66 h (in parallel to VEGF treatment) with: **A**, 0.2 µM nocodazole or 10 nM taxol; **B**, 10 µM blebbistatin or 0.5 nM calyculin A; **C**, 0.1 nM cytochalasin D + 0.1 nM latrunculin A or 150 µM CK666. Inverse aspect ratio data are averages from n=3 independent experiments, groups are compared using one-way ANOVA, Dunnett’s post-test (^∗^*P <* 0.05, ^∗∗^*P <* 0.01, ^*n*.*s*.^*P >* 0.05 represents statistical non-significance). Wrinkle depth data are pooled from n=3-7 independent experiments, groups are compared using Kruskal-Wallis, Dunn’s post-test (^∗∗∗^*P <* 0.001, ^*n*.*s*.^*P >* 0.05 represents statistical non-significance). Red arrowheads indicate the position of cell protrusions. Error bars represent ± SD. White scale bars, 50 µm. Yellow scale bars, 25 µm.

Surprisingly, all pharmacological treatments abrogating cell elongation also led to a significant increase in the depth of the out-of-plane displacements (fig. **8**), with the elongated wrinkles changing into a more dimply shape (fig. **8**A.b and (fig. **8**B.b)). Moreover, while inhibition of myosin light chain dephosphorylation with calyculin A resulted in deeper indentations, myosin inhibition with blebbistatin did not affect wrinkle depth (fig. **8**B.a). We thus hypothesized that the out-of-plane deformation could be promoted by cellular protrusive activity, a notion that was supported by observable protrusions under nocodazole (fig. **8**A.b) and calyculin A (fig. **8**B.b) treatment, as well as the fact that inhibition of actin polymerization with cytochalasin D and latrunculin A led to reduced wrinkle depth (fig. **8**C). To further test this hypothesis, we inhibited actin branching and lamellipodial formation using CK666, a specific inhibitor of the Arp2/3 complex (38). Arp2/3 inhibition resulted in increased wrinkle depth (fig. **8**C.a), suggesting that the potential implication of protrusions may involve filopodia rather than lamellipodia.

### Impaired cell elongation due to MT disruption induces the formation of vortex-like topological defects

Although nocodazole-induced MT disruption led to cell rounding as described in the previous section, sprouting events were not fully abrogated (an example can be observed in fig. **9**A). This provided an opportunity to investigate whether distortions of the director field were still involved in sprout initiation, ultimately reinforcing the notion of their importance in the process. Observation of cell shape and orientation revealed that the dimple regions were often associated with +1 topological defects (figs. **9**B and **9**C). Interestingly, the observed +1 topological defects exhibited a vortical or low-offset angle spiral nature, with cell orientations aligning nearly circumferentially around the center of the defect. The angle between the director field and the azimuthal direction from the center of the topological defect was − 3 ± 30º (*n* = 20 defects) at a radial distance of 75-100 µm from the center (fig. **9**B). Past theoretical work suggested that while aster- and spiral-like defects are unstable to out-of-plane deformations, vortex-like configurations should remain flat regardless of monolayer activity (39, 40). However, other studies have shown cell accumulation at the core of these defects (15, 41), potentially leading to out-of-plane deformations and possible tube formation (41). In line with this notion, our experimental measurements revealed a radial decay in local cell density that plateaus approximately 100 µm from the center of the defect (fig. **9**D). Moreover, the theoretical traction forces in eq. 1 around the dimple regions (fig. S8) can account for the observed dimple shape and depth (fig. **9**C), further supporting the relevance of distortions in nematic cell alignment for early angiogenesis.

**Fig. 9.**
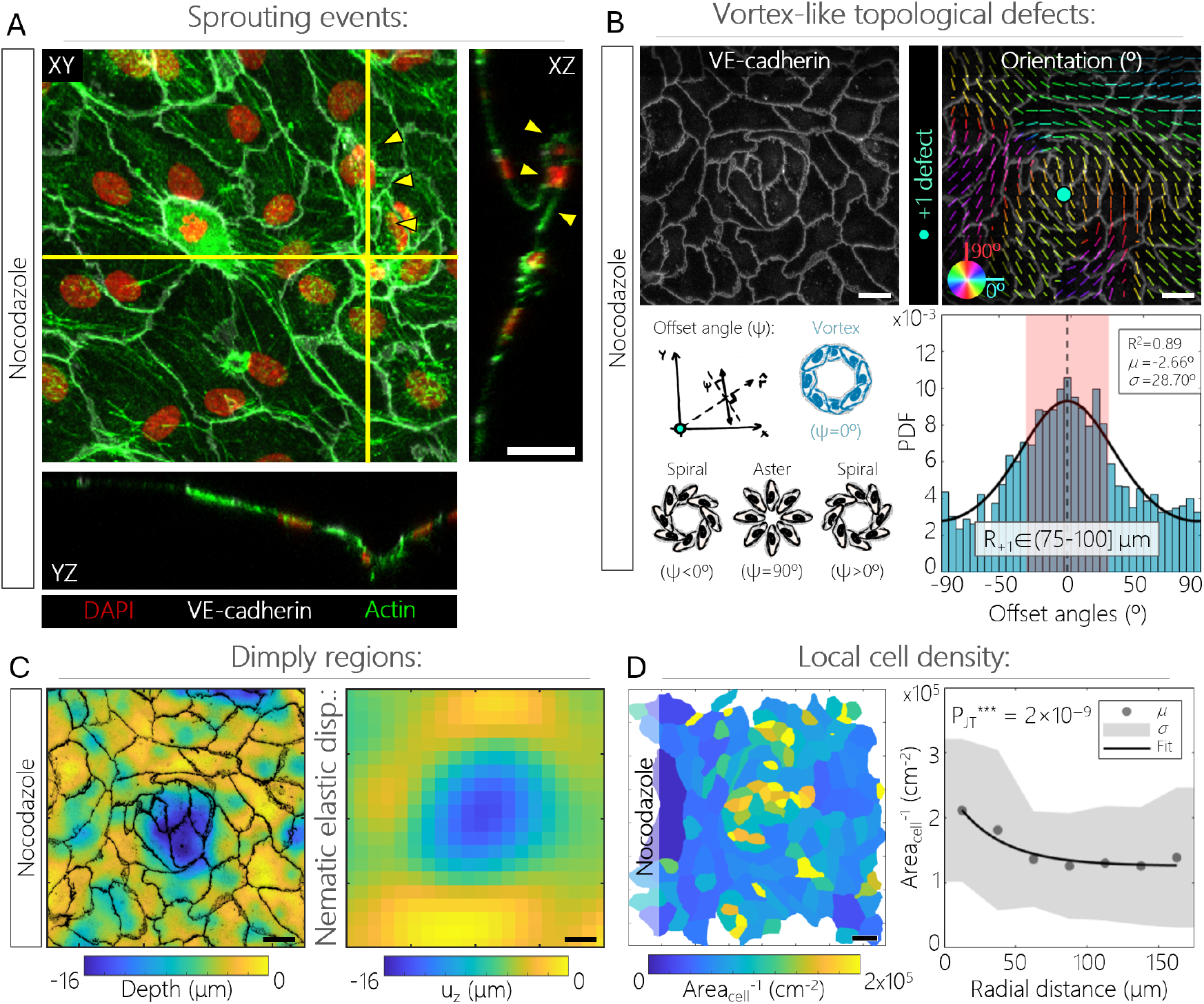
Nocodazole+VEGF treatment results in vortex-like defect formation. **A:** Immunostaining for nuclei (DAPI, red), actin (phalloidin, green) and cell-cell junctions (VE-cadherin, white) revealing a sprout (demarcated by yellow arrowheads). Scale bar, 25 µm. **B, top-left:** Maximum intensity projection of junctional immunostaining of an EC monolayer. **B, top-right:** Corresponding orientation map displaying a +1 topological defect (winding number +1). **B, bottom-left:** illustrated definition of the offset angle *ψ*: cell orientation refers to the tangential vector from the defect’s core; and possible +1 defects, their labels and offset angles. **B, bottom-right:** distribution of offset angles at 75-100 µm from the defects’ core. Pooled data (n=20 defects) from n=3 experiments. Black line: Von Mises distribution fit resulting in the *R*^2^ and mean (*µ*) values. The standard deviation (SD,*σ*, red shaded area) ignores the circular part (constant base) of the distributions. Scale bars, 25 µm. **C, left:** height map (VE-cadherin, black) showing a dimple at the level of the defect in **B, top**; **C, right:** reconstruction of the dimple height map observed in **C, left** as explained in Materials and methods. Scale bars, 25 µm. **D, left:** Colormap showing the inverse of the cell area around a vortex-like defect. **D, right:** Radial evolution of inverse of the cell area (proxy for cell density) from the defect’s center. The grey shaded area denotes the SD (*σ*). Pooled data (n=20) from n=3 experiments, Jonckheere-Terpstra test statistics for decaying trend (^∗∗∗^*P*_*JT*_ *<* 0.001). Black curve: exponential with an offset. Scale bars, 50 µm.

### Geometric and mechanical strategies for angiogenic control

Building on the results above, which suggest a significant role for distortions in nematic cell alignment in the initiation of angiogenic sprouting, we explored the potential of leveraging this finding for angiogenic control. Motivated by recent literature (13, 42, 14), we aimed to induce specific configurations of cell alignment by manipulating the boundary conditions at the edges of our cell monolayer. To achieve this, we resorted to the collagen hydrogel surface patterning method described in Materials and methods. With the aim of reducing the incidence of sprouting, we tested adhesive stripe patterns of different widths (400, 200 and 100 µm) to progressively vary the degree of cell alignment through physical confinement. The choice of pattern widths was based on preliminary experiments that pointed to an orientation correlation length on that order of magnitude for experiments either with or without VEGF (fig. S4B). While VEGF-treated HUVECs on 400 µm-wide adhesive stripes displayed weak alignment, the alignment progressively increased on 200 µm- and 100 µm-wide stripes, where cells consistently followed the main direction of the pattern, thus reducing the gradients in the nematic order parameter (fig. **10**A). The incidence of VEGF-mediated sprouting also decreased with decreasing pattern width, supporting the notion that distortions of the nematic field promote the occurrence of sprouting (fig. **10**A).

**Fig. 10.**
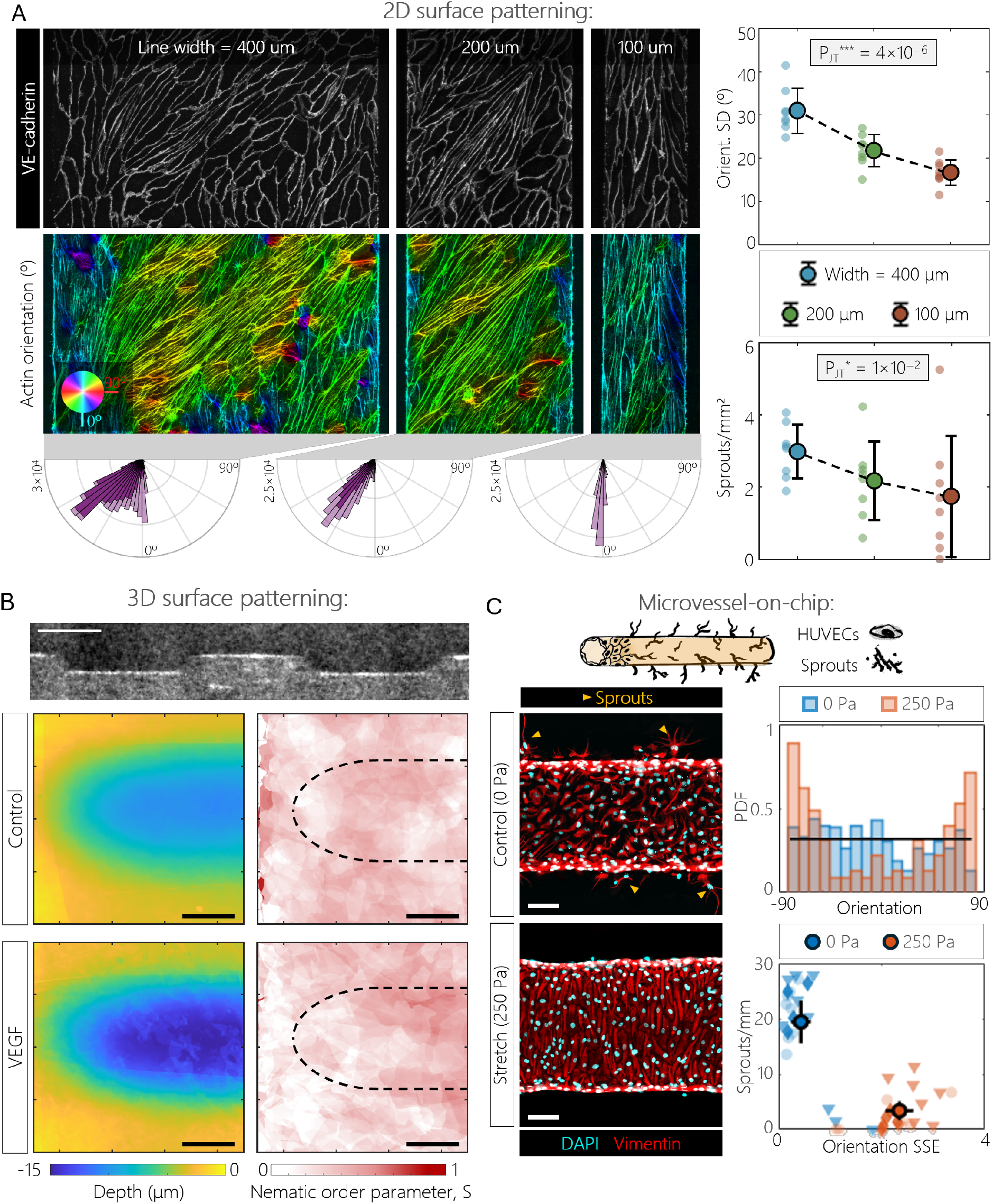
Biophysical strategies for angiogenic control. **A, left: 2D stripe patterns** – effect of pattern width (400, 200 and 100 µm). VE-Cadherin immunostaining (top, white) reveals the presence of cell streams, and phalloidin (bottom, F-actin) is color-coded by orientation angle. Histograms (pixel count) show overall orientation angle (light purple), and orientation angle away (*>*50 µm) from the pattern edges (dark purple). **A, right:** Spread (SD) of the actin orientation angle (top), and sprouting density (bottom) for the three widths. n=8 independent experiments, Jonckheere-Terpstra test statistics for trends (^∗^*P <* 0.05, ^∗∗∗^*P <* 0.001). Error bars represent ± SD. Discontinuous lines are guides for the eye. **B, left: 3D wrinkle patterns** – effect of surface topography. An OCT image of a cell-free pre-patterned collagen surface is followed by average height maps of cell-populated artificial wrinkles under control and VEGF conditioning. **B, right:** Average nematic order parameter shows no preferential cell alignment. Dashed curves approximate the artificial wrinkle. Scales bars 50 µm. **C, top: Microvessel-on-chip** – effect of pressure-induced stretch. Schematic of the cylindrical geometry. **C, bottom-left:** Immunostaining for vimentin (intermediate filaments, red) and DAPI (nucleus, cyan) of HUVECs under stretched (top) or control (bottom) conditions. Gold arrowheads demarcate sprouts. **C, right:** Top: Representative distributions of cell orientation angle under stretched (orange) and control (blue) conditions. Black line: uniform distribution. Bottom: Sprout density as a function of cell alignment (SSE: distance to a uniform distribution). Pooled data from n=3 independent experiments. Markers represent 1.3 mm long segments of the microvessel-on-chip, different symbols denote different experiments.

HUVEC alignment and orientation have also been shown to be controllable through substrate geometry at various scales (43–45). In this context, we wondered whether replicating the observed wrinkle geometry could trigger cell stream formation. Molding of the collagen surface prior to cell seeding produced 15 µm-deep, 100 µm-wide wrinkles. However, neither the control nor the VEGF-conditioned monolayers exhibited cell stream development at the level of these artificially induced wrinkles (fig. **10**B). We thus concluded that wrinkling was not the cause, but rather a consequence of cell stream formation.

The demonstration of angiogenic sprouting control by collagen hydrogel surface patterning as described above was conducted on a flat substrate. We explored if such control was also possible in a more physiologically relevant 3D setting. To this end, we resorted to an in-house developed EC-lined microvessel-on-chip in which we had previously demonstrated the ability to circumferentially align the cells in response to pressure-induced stretch (46, 47). In this system, stretch indeed resulted in increased EC alignment, which, importantly, was accompanied by significantly reduced sprouting, in line with our hypothesis that distortions in cell nematic order favor the initiation of angiogenic sprouting.(fig. **10**C). These findings provide further support for the potential of exploiting EC topological modulation for angiogenesis control.

## Discussion

The process of sprouting angiogenesis, whereby new blood vessels emerge from pre-existing vasculature, is of fundamental interest in diverse fields such as tumorigenesis (48), vascular pathologies (49), and tissue regeneration (50). While previous studies have documented the role of VEGF, a potent pro-angiogenic factor, in orchestrating EC shape changes and modulating cell proliferation and migration, a major challenge remains in elucidating how these diverse cellular activities combine to drive specific morphogenetic events. In the present study, we aimed to address this gap by investigating how EC organization contributes to angiogenic sprouting. Specifically, we explored how the induction of nematic domains by VEGF conditioning results in mechanical forces and substrate deformation, influencing the initiation of sprouts.

Treating HUVEC monolayers cultured on a fibronectin-coated collagen I hydrogel substrate with VEGF induced an increase in average cell elongation, accompanied by spatial heterogeneity in cell morphology. This heterogeneity was characterized by the emergence of elongated, aligned cell streams that coexisted with more polygonal cell regions. These observations are consistent with previous studies that have provided qualitative evidence for the influence of VEGF on endothelial morphology. For instance, Vion et al. observed a significant increase in cell elongation when ECs were treated with VEGF-A (21), and Cao et al. reported that VEGF stimulation in confluent EC monolayers prompted gradual elongation in only a subset of the cells (20). Interestingly, angiogenic sprouts preferentially originated from the tips of these highly ordered domains, regions that we termed “wedges”.

Modeling our substrate as a linear elastic semi-infinite space with a Poisson’s ratio between 0 and 0.5, we showed that the observed forces could induce both in-plane and out-of-plane displacements in the collagen substrate, leading to the formation of micron-scale wrinkles. Experimentally, ∼ 4–5 µm-deep wrinkles were in fact observed with VEGF treatment and were localized under the elongated cell streams. However, while our model predicted that out-of-plane deformations should be at least 2 to 3 times smaller than in-plane displacements, we observed similar magnitudes in both dimensions. Additionally, our model predicted that wrinkle formation should be localized at the wedge, but our experimental results demonstrated that wrinkles extended along the entire cell stream. Although these discrepancies may be partially attributed to the complex mechanical properties of collagen, including factors such as plasticity, anisotropy, and collagen hydrogel response to multi-axial loads (30–32, 51), the differences may not be due to material properties alone. The observed deformation patterns may also be influenced by additional mechanisms such as extracellular matrix (ECM) remodeling. However, although VEGF has previously been shown to upregulate the deposition of BM components including fibronectin and collagen IV (52), we observed no significant spatial heterogeneities in BM protein deposition that could explain the observation of wrinkle formation. Although previous studies have shown that VEGF can boost the production of matrix-degrading enzymes of the metalloproteinase family (53, 54) and although we observed slightly increased levels of MT1-MMP within the cell streams, broad spectrum inhibition of MMPs did not reduce wrinkle formation. MT1-MMP has been shown to also regulate VEGF-A expression (55), which suggests that its role in angiogenesis may go beyond matrix degradation. Given this, the slight increase in MT1-MMP levels within the cell streams observed in our study might reflect additional, non-degradative functions, further complicating the potential role of matrix degradation in wrinkle formation.

Our findings also indicate that cells along the stream direction experience tensile or compressive stresses depending on their position relative to the wedge. Previous studies show that cells under tensional states (inside the streams in our case) experience upregulation of VEGFR-2 expression (56) and phosphorylation (57), with this receptor being indispensable for VEGF-induced cell elongation (20), suggesting a positive feedback loop since VEGF-A primarily interacts with ECs through VEGFR2. VEGF is known to promote cell polarization by influencing MT organization (58) and to modulate actomyosin contractility by regulating actin filament polymerization and depolymerization (59) as well as promoting myosin light chain phosphorylation (60). Consistent with these observations, pharmacological perturbations revealed that MT dynamics are crucial for cell stream formation, with MT polymerization disruption by nocodazole or stabilization by taxol impairing cell elongation. Additionally, we showed that actomyosin contractility counteracts cell elongation. More specifically, increased contractility with calyculin A promoted a cuboidal cell morphology, while myosin inhibition with blebbistatin and actin depolymerization using cytochalasin D and latrunculin A slightly enhanced overall elongation.

Interestingly, while cytoskeletal perturbation significantly altered cell morphology and stream formation in a predictable manner, it had some unexpected effects on the out-of-plane deformations. Given that actomyosin contractility is widely viewed as the primary driver of traction forces, we anticipated that blebbistatin would reduce wrinkle depth, but this was not the case. It should be noted that while the role of non-muscle myosin II is well established (61, 62), the body of literature reporting the effects of blebbistatin on endothelial contractility is very small, with one relevant study showing the need for combining blebbistatin with Y27632 to induce notable reductions in cell-exerted traction forces (56), which may explain our current findings. In contrast, hampering actin polymerization with cytochalasin D and latrunculin A did result in reduced wrinkle depth, supporting the hypothesis that contractile forces exerted by the actomyosin machinery play an important role in the observed substrate deformations. Moreover, all three pharmacological treatments resulting in cell rounding (nocodazole, taxol and calyculin A) led to the formation of dimples of increased depth instead of wrinkles. Under nocodazole conditioning, the dimples correlated with the appearance of vortex-like arrangements in the EC monolayer, which is consistent with the traction forces derived from our nematic model, further supporting the role of nematic alignment in substrate deformation. We also hypothesized that protrusive activity may have a major influence on wrinkle formation. CK666 treatment, which inhibits Arp2/3 branching and lamellipodial formation, resulted in increased wrinkle depth, pointing to a complex but significant role of protrusive activity in 3D deformation, most likely via filopodia which indeed are the favored form of protrusions in the absence of Arp2/3 nucleators (63).

Intriguingly, despite the disruption of cell elongation induced by nocodazole treatment, angiogenic sprouts were still observed to emerge at sites exhibiting increased out-of-plane deformation. This finding prompted us to explore whether these locations exhibited specific topological characteristics independent of cell elongation. Our analysis revealed that the combined treatment of VEGF and nocodazole induced ECs to form vortex-like topological defects. Consistent with recent studies, these defects displayed local accumulation of cell density at the core of the defect (15, 41). This increase in cell density can, in turn, force an out-of-plane deformation whose shape depends on the fluidity of the tissue. Interestingly, the theory shows that cell-neighbor exchange can lead to the development of tube-like structures (41).

The findings described above indicate that angiogenic sprouting initiates preferentially at sites with elevated gradients in the nematic order parameter, suggesting that modulating cell alignment and elongation can potentially serve as a means to control sprouting. We investigated this hypothesis in both 2D and 3D systems. In 2D, we reinforced cell alignment and thus reduced gradients in nematic order by patterning the collagen hydrogel surface with adhesive stripes on a scale of the order of the system’s correlation length. Previous reports have shown that ECs and other cell types reliably follow the edges between adhesive and non-adhesive zones (13, 14, 64). Indeed, the increase in cell alignment generated by the collagen surface patterning technique developed here resulted in a reduction in sprout initiation. In 3D, as an initial step toward elucidating the role of topology in mediating the angiogenic response to mechanical cues, we demonstrated that static stretch suppresses sprouting by promoting circumferential cell alignment in an EC-lined microvessel-on-chip platform. It is worth noting that the effects of static strain on angiogenic sprouting have been largely overlooked. Although one study reported pro-angiogenic effects (65), it unfortunately did not include direct observations of cell morphology or alignment.

While our findings provide strong evidence for the role of cellular organization in sprout initiation, several aspects remain to be explored. First, although we identified a clear relationship between local nematic order gradients and sprouting as well as between local nematic order gradients and substrate deformation, we did not directly quantify whether wrinkle depth correlates with sprouting frequency. Future studies should investigate this relationship to determine whether stress or strain actively contributes to sprout emergence. Secondly, while our TFM experiments provide valuable static snapshots of force distribution, they lack the temporal resolution to capture dynamic changes in cell traction during sprouting. Angiogenic sprouting is a time-dependent process, and real-time measurements of traction forces promise to provide insight into the role of forces at different stages of cell stream and sprout formation. Additionally, performing 3D TFM would help determine whether protrusive activity is directly linked to these processes. It is also important to consider how our experiments relate to *in vivo* systems. At scales comparable to the wrinkle sizes observed in our experiments, high VEGF levels in the absence of VEGF gradients have been shown to promote vessel enlargement (66–68), a phenomenon that bears some resemblance to the wrinkling process. Although often linked to cell proliferation, the exact mechanism behind this luminal expansion remains unclear and may be related to the observed 3D deformations.

The current study focused exclusively on VEGF as an angiogenic sprout-promoting agent. The role of other pro-angiogenic factors such as fibroblast growth factor (FGF) and platelet-derived growth factor (PDGF) remains to be explored, and it would be particularly interesting to determine whether these other molecules also rely on active nematic traction forces to initiate sprouting. Furthermore, our study did not explicitly address the role of junctional dynamics in the sprouting process, even though the collective behavior of EC monolayers during sprouting angiogenesis is known to be dictated by both cytoskeletal dynamics and cell-cell interactions. Lastly, while cytoskeleton-perturbing drugs modulated cell behavior in our experiments, incomplete disruption of cytoskeletal components may have influenced the observed effects, as compensatory mechanisms may have possibly mitigated some of the expected changes.

Taken together, our findings provide strong evidence that gradients in EC nematic order serve as predictive markers for sprouting angiogenesis. We propose that nematic order, active traction forces, and VEGF signaling interact to regulate angiogenic sprout initiation. These insights open exciting avenues for mechanobiological control of angiogenesis, with potential applications in tissue engineering, regenerative medicine, and anti-angiogenic therapies. Future research should focus on quantifying the interplay between pro/anti-angiogenic signaling and nematic features, investigating how mechanical constraints influence vascular morphogenesis, and developing geometric control strategies to direct angiogenesis in engineered tissues.

## Materials and methods

### Fabrication of experimental platforms

#### Collagen hydrogel preparation and attachment

Hydrogels of type I rat tail collagen with a pH of 8 (at 37 ^°^C and 5 % CO_2_) and a concentration of 4 mg*/*mL were used for all experiments. The detailed protocol used for hydrogel fabrication can be found elsewhere (69). Briefly, collagen I was isolated from rat tail tendon using HCl, resulting in an acidic solution. This solution was then neutralized with a NaOH buffer. Collagen hydrogel polymerization was achieved in a tissue culture incubator for 1-2 h. To improve collagen hydrogel attachment to the hydrophobic PDMS wells in which it was cast, the PDMS surface was plasma-activated and coated first with 1 % polyethylenimine (Sigma-Aldrich, Burlington, MA, USA) in H_2_O for 20 min and then with 0.01 % glutaraldehyde (Sigma-Aldrich) in H_2_O for 20 min before final rinsing with phosphate buffered saline (PBS). Polyethylenimine treatment was combined with UV irradiation for sterilization purposes.

#### Flat collagen hydrogel system

A PDMS stencil (well diameter: 4-5 mm, thickness: 250 µm; Alveole, Paris, France) and a round coverslip (coverslip diameter: 18 mm; Epredia Menzel-Gläser, Kalamazoo, MI, USA) were bonded together using plasma activation. Collagen hydrogel was then cast into each of the wells, and its free surface was maintained flat by covering it with a second coverslip that was subsequently removed after hydrogel polymerization. The hydrogel surface was coated for 1 h with fibronectin (Sigma-Aldrich) at a concentration of 50 µg*/*mL in PBS. Samples were inserted in a 12-well plate (Corning Inc., Corning, NY, USA) for subsequent experiments.

#### Adhesive/non-adhesive surface patterning

4-arm poly(ethylene glycol)-acrylate (10 kDa; Laysan Bio Inc., Arab, AL, USA) was crosslinked to the collagen hydrogel surface to hamper fibronectin and cell attachment following a prescribed pattern. Micropatterning was performed with a PRIMO patterning system (Alveole). The process, schematized in fig. S9, is as follows: a thin polyethylene (PE) film is laid on the inner part of a 35 mm-diameter Fluorodish (World Precision Instruments Inc., Sarasota, FL, USA). A mixture of 15-30 µL of PLPP photoinitiator (Alveole) and 4-arm PEG acrylate at a concentration of 50 mg*/*mL is then deposited on the plastic film and squeezed by placing the sample face-down (note: lifting and lowering the sample several times ensures penetration of the mixture into the sample). UV illumination is then applied with focus at the interface between the PE and the hydrogel, following patterns stored in grayscale 8-bit images (TIFF format), with illumination being proportional to pixel brightness. A UV dose of 600 mJ*/*mm^2^ is necessary to obtain clean non-adhesive regions.

#### Topographical surface patterning

A PDMS template with bumps of the desired shape was laid on top of the system during collagen polymerization, imprinting concave shapes (the negative of the template) in it. As shown in fig. S10, the steps leading to the final PDMS template include the polymerization of a 15 µm-thick dry film photoresist (Eternal Materials Co., China) using the PRIMO system. The photoresist was first attached to a glass coverslip with the help of a speed-controlled office laminating machine heating up to 120 ^°^C (PEAK®Performance PS-320, Vivid Laminating Technologies, United Kingdom). The desired pattern was then used for UV illumination at a very low dose (7-8 mJ*/*mm^2^). After washing off the unpolymerized film with a 5 % solution of potassium carbonate (Sigma-Aldrich) in H_2_O, two PDMS-casting stages were implemented, resulting in first a negative and finally a positive PDMS template. To enable glass-PDMS and PDMS-PDMS casting, the master molds were silanized by plasma activation and exposure to trichloro(1H,1H,2H,2H-perfluorooctyl)silane (Sigma-Aldrich) vapor for 20 min.

#### Microvessel-on-chip

The microvessel-on-chip system (fig. S11) was fabricated as described previously (46). Briefly, in a PDMS chamber, a hollow channel was created within a collagen hydrogel by casting a 350 µm-thick needle. The needle was then removed, and cells were pipetted into the channel. Flow in the microvessel was imposed using a syringe pump (Harvard Apparatus). An outlet reservoir imposed the pressure in the channel, while the inlet reservoir ensured delivery of the imposed flow rate. The open top of the system allowed pressure in the channel to generate circumferential strain.

#### TFM collagen substrates

The collagen substrates used in the TFM experiments were identical to the flat collagen hydrogel system described above with one exception: 0.5 µm-diameter fluorescent beads (17152-10, Fluoresbrite) were embedded in the collagen hydrogel at concentrations ranging from 0.00625 to 0.025 % (v/v).

### Cell culture

Human umbilical vein ECs (HUVECs; ScienCell Research Laboratories Inc., Carlsbad, CA, USA) were cultured in Endothelial Cell Medium (ECM) containing 5 % fetal bovine serum, 1 % EC growth supplement and 1 % penicillin-streptomycin (ScienCell) at 37 ^°^C in a humidified atmosphere with 5 % CO_2_. At early confluence, cells were trypsinized with TrypLE enzymes (Gibco, Thermo Fisher Scientific) and seeded in the experimental platforms. A seeding density of ∼ 50 000 cells*/*cm^2^ was used in all experiments except the microvessel-on-chip where seeding was performed at an already confluent density, as detailed elsewhere (46). Cells were used between passages 3 and 6, with splitting ratios below 1:10.

### VEGF conditioning

48 h after seeding, ECs were subjected to 50 ng*/*mL VEGF for 66 h. Control experiments were performed in the absence of this growth factor.

### Pharmacological treatment

#### Cytoskeletal disruption

The cytoskeletal-altering drugs used included: nocodazole (M1404, Sigma-Aldrich) at 0.2 µM, paclitaxel/taxol (89806, PhytoLab) at 10 µM, blebbistatin (B0560, Sigma-Aldrich) at 10 µM, Y-27632 dihydrochloride (Y0503, Sigma-Aldrich) at 10 µM, calyculin A (208851, Sigma-Aldrich) at 0.5 µM, cytochalasin D (C2618, Sigma-Aldrich) at 20 nM, and latrunculin A (428026, Millipore) at 10 nM. All reagents were incubated for 66 h in parallel to VEGF conditioning. An initial study on the effects of DMSO concentration showed no significant effect on the measured cell inverse aspect ratios and wrinkle depths (fig. S12).

#### Inhibition of matrix metalloproteinases

Ilomastat (M5939, Sigma-Aldrich), a broad-spectrum MMP inhibitor, was used at 10 µM simultaneously with VEGF.

### Microvessel perfusion and pressure application

ECs in the microvessel were cultured to confluence under low flow conditions (2 µL*/*min) for the first 24 h. The same flow rate was then maintained for another 24 h under either low pressure (outlet pressure=0 Pa) or high pressure (outlet pressure=250 Pa) conditions to induce circumferential stretch in the EC monolayer. Outlet and inlet reservoirs were used as previously explained (46) to provide the desired hydrostatic pressure difference and mean level for the targeted steady flow and pressure.

### Immunostaining

At the end of each experiment, samples were fixed in 4 % paraformaldehyde (Thermo Fisher) in PBS for 15 min and were subsequently rinsed with PBS. Cell permeabilization with 0.25 % Triton® X-100 solution in PBS was then performed for 30 min, followed by saturation with 3 % bovine serum albumin (BSA) in PBS for 1 h. Samples were then incubated for 1 h in a solution of primary antibodies in PBS. Primary antibodies were used at 1/400-1/200 dilutions and included: rabbit anti-VE-cadherin (ab33168, Abcam), mouse anti-ZO-1 (33-9100, Thermo Fisher), mouse anti-*β*-catenin (C7082, Sigma-Aldrich), mouse anti *α*-tubulin (Sigma-Aldrich, T5168), rabbit anti-TGN46 (ab50595, Abcam), mouse anti-CD34 (MA1-10202), mouse anti-fibronectin (ab6328, Abcam), rabbit anti-collagen IV (ab6586, Abcam), rabbit anti-laminin (ab11575, Abcam), mouse anti-MMP14 (MAB3328), apt for HUVECs. After rinsing 3 times in PBS for 10 min, a final incubation step was applied including 4’,6-diamidino-2-fenilindol (DAPI), phalloidin Alexa fluor 488 (Life Technologies) at 1/400 dilution and the following secondary antibodies: Alexa fluor 555-conjugated donkey anti-rabbit antibody (ab150074, Abcam) and Alexa fluor 647-conjugated goat anti-mouse antibody (ab150115, Abcam), or Alexa fluor 555-conjugated donkey anti-mouse antibody (ab150106, Abcam) and Alexa fluor 647-conjugated donkey anti-rabbit antibody (ab150075, Abcam) at 1/400-1/200 dilution. A long final rinsing (over 24 h, maximizing PBS exchanges) appears to be best at reducing background noise. The entire immunostaining protocol was performed at room temperature.

### Microscopy

#### Epifluorescence microscopy

Stack acquisitions of immunostained samples were obtained using an inverted microscope (Nikon Eclipse Ti) with a 10X objective (CFI Plan Fluor DLL 10X, MRH10101), imposing vertical steps of 5-10 µm.

#### Confocal microscopy

Stack acquisitions of fixed samples were obtained on an inverted confocal microscope (Leica TCS SP8) using a 40X objective (HC PL APO 40x/1,1 W CORR CS2, 11506425), with z-steps of 0.25-0.5 µm.

#### Traction force microscopy

The polystyrene fluorescent beads embedded in the collagen hydrogel were imaged at two time points for each experiment: right before seeding (reference state), and after fixation (stressed state) at the end of the experiment (66 h under the desired conditioning). 0.5-1 µm-spaced z-stacks containing the beads of interest were acquired using a spinning disk confocal microscope (X-Light, CrestOptics) with a 40 µm pinhole size and a 20X objective (CFI Plan Fluor DLL 20X, MRH10201). Because it is impossible to predict the location of cell streams *a priori*, several adjacent (and slightly overlapping) fields of view were acquired every time to increase the probability of finding cell streams at the last time point.

#### Line-field confocal optical coherence tomography

In the absence of a cell monolayer, we turned to a LC-OCT microscope (Damae Medical) to perform topography measurements. 3D acquisitions resulted in 1230×500×700 µm volumes, with a voxel resolution of 1 µm in all three axes.

### Data analysis

#### Sprouting quantification

Sprout identification in widefield stacks was performed based on their appearance (similar to confocal) and under the condition of their constitutive actin/VE-cadherin bundles spanning over 10 µm in the direction perpendicular to the monolayer (out-of-plane). Sprout density analysis was performed, and sprout coordinates were stored using an in-house developed MATLAB code.

#### Cell morphology analysis

Cell segmentation in EC monolayers was automatically performed using Cellpose (70) on immunofluorescence images of cell-cell junctions (*β*-catenin, ZO-1, or VE-cadherin). Based on *Cyto 2*, we trained our own model to improve the segmentation of highly elongated cells. On the basis of this segmentation, the inverse aspect ratio of each cell, defined as the ratio of the minor to major axis of a fitted ellipse, was calculated using MATLAB.

#### Detection of cell streams, wedges and distance computation

A Cellpose model was trained to segment polygonal cells only (a comparison to a thresholding approach is provided in fig. S13A). The resulting images were binarized (with cell stream regions containing 1’s and polygonal cell regions 0’s) and processed in MATLAB to obtain differentiable regional boundaries. In short, after small regions were removed by thresholding, the remaining regional edges were extracted and underwent a moving average processing step. Wedges were then defined as the regions where the edge of a cell stream was perpendicular to the cells in it. Using a custom-made MATLAB code, cell orientation (*θ*_*cells*_) was estimated from the brightness gradients of cell-cell junction immunostaining images, and regional boundary orientation (*θ*_*boundary*_) from its extracted binary shape. In computational terms, perpendicularity was defined as:

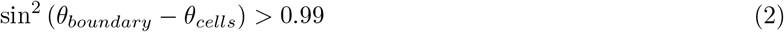

Distance maps to wedges could then be calculated and values extracted for the pixels in cell streams, polygonal cell areas, and those corresponding to sprout coordinates. Each region was treated separately, and histograms were calculated pooling all data.

#### Assessment of the p-atic orientational order

The magnitude of the p-atic orientational order as a function of the coarse-graining radius was computed following the method described by Armengol-Collado et al. (71), using the code provided in their associated GitHub repository. Briefly, Armengol-Collado et al. introduced a rank-p shape tensor designed to capture arbitrary p-fold rotational symmetries in polygonal cell contours, where *p* is any natural number. Each component of this tensor is proportional to a characteristic ‘shape function’ that depends on the geometry of the polygon. For a cell with *V* vertices located at positions *r*_*v*_ relative to the cell center, and angles *ϕ*_*v*_ measured between each vertex and the x-axis, the shape function is given by:

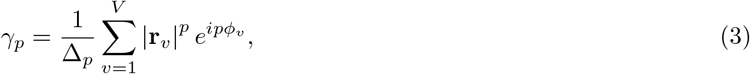

This shape function can be coarse-grained, enabling quantification of liquid crystal order. The magnitude of the resulting coarse-grained parameter |Γ_*p*_ |, shown in fig. S4A for both control and VEGF-conditioned monolayers, serves as a measure of orientational coherence within the specified coarse-graining radius.

#### Assessment of the director field

The director field **n** was calculated cell-wise. Briefly, an ellipse was fitted to the boundary of the cell, and the coordinates of its major and minor axes were extracted. As the director field remains invariant under the transformation **n** → − **n**, to avoid ambiguities, all the subsequent operations were performed on the components of the Q-tensor, which was calculated as:

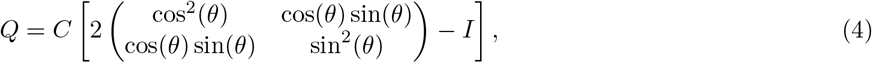

where *θ* is the orientation of the major axis with respect to an arbitrary x-axis and *C* = (*a* − *b*)*/*(*a* + *b*), with *a* and *b* the major and minor axes of the cell, respectively. The components of the Q-tensor were then averaged over different images and over a disk of radius R_coarsening_=1 µm for figs. **4**A and S8A and R_coarsening_=40 µm for figs. S5 and S8B and all nematic traction force assessments.

The largest eigenvalue, *λ*_*max*_, of the coarsegrained Q-tensor corresponds to the scalar nematic order parameter, and its principal eigenvector, **V**_*principal*_, to the orientation of the director field **n**. Finally, we define the director field **n** as:

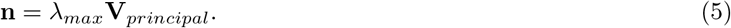

#### Computation of the nematic traction forces

Following Refs. (25, 15, 26), the nematic traction forces generated by activity can be expressed as in eq. 1. Note that the three force terms in this equation depend on the director field and its gradients only up to a constant material parameter: *t*_*modulus*_, *t*_*splay*_, and *t*_*bend*_.

Using the director field **n**, which was computed by the method above, we determine each of the contributions to the nematic traction forces in eq. 1. Specifically, fig. S6 shows each of those contributions separately for two experimental cases: control EC monolayers and VEGF-treated EC monolayers. Similarly, fig. S8B shows each of those contributions separately for EC monolayers treated with VEGF and Nocodazole.

Fig. **4**C shows the traction force map computed for the contributions in eq. 1 of VEGF-treated EC monolayers (fig. S6) and the values of the material parameters: *t*_*modulus*_ = 16 Pa · µm, *t*_*splay*_ = 227 Pa· µm, *t*_*bend*_ = − 106 Pa ·µm. Figs. **4**C and **5**A show the in-plane and out-of-plane displacement fields on a half-space linear elastic material subjected to such traction forces. Finally, fig. **9**C shows the z-displacement field on a half-space linear elastic material that is produced by a traction force map computed for the contributions of eq. 1 in fig. S8B and the values of the material parameters: *t*_*modulus*_ = −16 421 Pa·µm, *t*_*splay*_ = −44 356 Pa·µm, *t*_*bend*_ = 61 578 Pa·µm.

#### 2D Traction force microscopy

TFM analysis was preceded by average projection of the image stacks containing the fluorescent beads in the first ∼ 25-30 µm from the surface. 2D PIV was then performed using PIVlab (72), using the relaxed state (before cell seeding) as a reference. Cell-substrate tractions were then calculated using the Fourier transform traction microscopy suite proposed in Saraswathibhatla et al. (73), with E_substrate_= 3 × 10^2^ Pa, *ν*_substrate_=0.3, and h_substrate_=250 µm.

Displacements and tractions around wedges were finally averaged after alignment of all fields of view based on the wedge position and the average cell orientation in the cell stream close to the wedge, as obtained from cell-cell junction immunostaining (fig. S14).

#### Gaussian load on a half-space linear elastic material

Here, we derive the equations for the displacement field generated by a Gaussian load that is applied on the boundary of a half-space linear elastic material.

The collagen substrate is described as a linear elastic material in a half-space geometry *z* ≤ 0 with a plane in contact with the external medium (i.e. cell monolayer) at *z* = 0. The displacement field **u** describes the deformation of a material point. The strain tensor is *ϵ*_*ij*_ = (*∂*_*i*_*u*_*j*_ + *∂*_*j*_*u*_*i*_)*/*2, where the indices *i* and *j* run over the three Cartesian directions (*x, y, z*). For a linear elastic material, the stress field is related to the strain tensor by the expression:

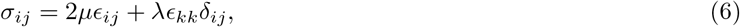

where *δ*_*ij*_ is the identity matrix, and the elastic parameters *µ* = *E/*(2(1 + *ν*)) and *λ* = *Eν/*((1 + *ν*)(1 − 2*ν*)) are related to the Young’s modulus *E* and the Poisson’s ratio *ν*. At steady state, the momentum conservation reduces to a force balance:

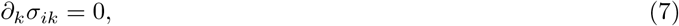

On the upper boundary *z* = 0, we enforce the force balance

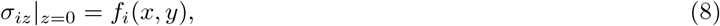

where *i* runs over the Cartesian indices (*x, y*) and **f** (*x, y*) = (*f*_*x*_(*x, y*), *f*_*y*_(*x, y*)) represents the traction force density generated by a cell or a group of cells on the collagen substrate. Here, we consider that this force density is centered at *x* = *x*_0_ = 0 and *y* = *y*_0_ = 0, and has a Gaussian distribution:

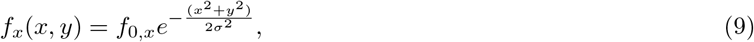

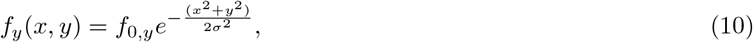

where *f*_0,*x*_ and *f*_0,*y*_ are the values of the force density components at *x* = 0 and *y* = 0 and the parameter *σ* quantifies the spread of the force density. Finally, on the bottom boundary 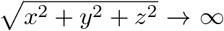, we enforce that the displacement field vanishes.

At *z* = 0, the displacement field that satisfies the model above is given by:

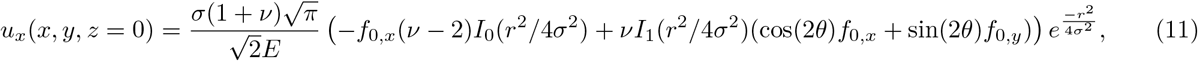

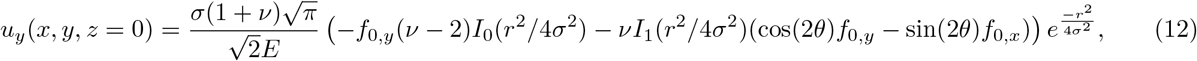

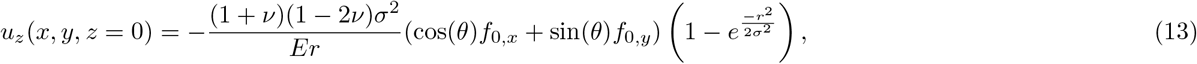

where 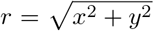 and *θ* = *Arg*(*x* + *iy*). Note that in the incompressible case where *ν* = 1*/*2, the force density **f** (*x, y*) does not generate an out-of-plane displacement, i.e. *u*_*z*_(*x, y, z* = 0) = 0. Finally, we note that the net force exerted by the cell monolayer on the substrate must vanish.

To produce the displacement fields in figs. **4**C and **5**A equations eq. 11, eq. 12, and eq. 13 were used for a distribution of Gaussian loads that matched the traction force map obtained by TFM in fig. **4**B. To produce the *z*-displacement field *u*_*z*_(*x, y, z* = 0) in fig. **9**C equation eq. 13 was used for a distribution of Gaussian loads that matched the experimental height map in fig. **9**C. Unless stated otherwise, the Young modulus was *E* = 300 Pa and the Poisson’s Ratio was *ν* = 0.3.

#### Wrinkle depth analysis

For monolayer confocal stacks, height maps were derived from the brightest z position in the phalloidin channel. After Gaussian smoothing, the linear component (corresponding to tilt or large undulations of the gel) of the surface was removed, and wrinkle depth was calculated as the difference between the highest and lowest coordinates in the map. The size of all acquired images was 290×290 µm (∼ 3 times the width of a cell stream), with a resolution of 0.57 µm*/*px. Heightmaps containing wedges were aligned as previously explained (fig. S14). In LC-OCT images, surface height maps were obtained by fitting step or Gaussian functions to the vertical intensity profiles. Successive wrinkle depth analyses were performed as explained above.

#### Basement membrane thickness quantification

Confocal stacks containing fibronectin, collagen IV or laminin images were analyzed, and thickness maps were constructed by measuring the full width at half maximum (FWHM) of a Gaussian fit of the z intensity profile at each XY coordinate. Cell stream and polygonal cell zones were classified, and region-wise average thickness was calculated and compared to the overall average thickness in each image.

#### MT1-MMP regional quantification

MT1-MMP intensity was calculated as 8-bit pixel values normalized by 2 × 10^8^. Cell stream and polygonal cell zones were classified, and region-wise average intensity was calculated and compared to the overall average intensity in each image.

#### Actin and microtubule orientation analysis

Cytoskeletal orientation was calculated from intensity gradients revealed by phalloidin and *α*-tubulin staining in confocal images. This operation was performed using either the FIJI plugin OrientationJ (74, 75) or an in-house developed MATLAB function based on the same references.

#### Detection of +1 topological defects and classification

Orientation maps were calculated using OrientationJ on cell-cell junction (VE-cadherin) immunostaining images, then plugged into a defect detection MATLAB code. The detection algorithm is based on the calculation of the winding number. Offset angles were calculated as the angles between local orientation and the local tangential vector from the defect’s core. Offset angles were averaged over different radii from the center of the topological defect (R=0-25 µm, 25-50 µm, … 150-175 µm). Offset values close to 0^°^ indicate the presence of a vortex, while increasing values denote a spiral character, and ultimately an aster (90^°^).

#### Quantification of cell density profiles around vortex defects

Cells were segmented and the inverse of the resulting cell area was averaged over different radii similarly to the previous epigraph.

#### Sprouting and cell alignment in the microvessel-on-chip

Linear sprout density was manually quantified considering the number of structures visible outside the edge of the cylindrical microvessel on a side view of vimentin immunostaining. Cells inside the microvessel (top or bottom half) were segmented using Cellpose’s *Cyto 2* model on the vimentin staining, the cylindrical surface was developed for curvature correction, and cell orientation distributions were calculated by ellipse fitting with MATLAB. The sum of squared errors (SSE) relative to a uniform distribution was used as a measure of the extent of cell alignment.

#### Other third-party codes

The Stack Focuser plugin in FIJI (74, 76) and MATLAB’s *fstack* function (77) were used to focus epifluorescence and confocal images. Ellipse fitting to cell contours was performed in MATLAB using the *planarFit* function (78) and the *MOMENT* function (79). Jonckheere-Terpstra statistical tests were performed in MATLAB using the *jttrend* function (80).

### Statistical analysis

Statistical tests were performed using MATLAB. All statistical analyses in the main text were based on three or more independent experiments. Sprout densities were compared using a two-sample t-test on the distributions of per-experiment averages (fig. **1**C). Distributions of per-experiment average cell inverse aspect ratios in figs. **2**B and **8** were all compared together using a one-way ANOVA followed by a Dunnett’s post-test, with VEGF used as the control group (averages are assumed to be normally distributed). Wrinkle depth data were pooled for each condition and compared using Kruskal-Wallis followed by Dunn’s post-test to avoid normality assumptions. Wrinkle depth data derived from confocal images (figs. **6**C and **8**) were compared together, separately from data derived from LC-OCT images (fig. **5**C). Analyses of regional BM thickness (fig. **6**A) and regional MT1-MMP intensity (fig. **6**B) were performed on pooled data using a two-sided Wilcoxon rank sum test. Trends in cell density around vortex-like defects (fig. **9**D) and in the SD of orientation and sprout density in stripe patterns (fig. **10**A) were analyzed using a Jonckheere-Terpstra test.

Statistical analyses in the supplementary materials were performed separately, as the analyzed data were extracted from experiments other than those in the main text. The selected statistical tests were the same as in the main text: one-way ANOVA followed by a Dunnett’s post-test for average cell inverse aspect ratios (fig. S12), Kruskal-Wallis followed by Dunn’s post-test on pooled data of wrinkle depth (fig. S12), and two-sided Wilcoxon rank sum tests on data of regional BM thickness (fig. S7). Exceptionally, supplementary data on regional BM thickness correspond to a single experiment for each condition (fig. S7).

Details regarding the number of data points per experiment, the specific statistical tests used, and the significance levels are provided in the corresponding figure legends.

## Data and code availability

Data and codes are available upon request to the corresponding author.

## Acknowledgments

We are grateful to Dr. Pau Guillamat for generously sharing the MATLAB code used in our defect detection analysis. We thank Dr. Charlie Duclut and Professor Jacques Prost for fruitful discussions. We also acknowledge Dr. Aurélien Duboin (Alvéole) for assistance with patterning experiments, and Dr. Gaël Latour, Dr. Pierre Mahou, and the Laboratory of Optics and Biosciences (LOB) at École Polytechnique for their support with line-field confocal optical coherence tomography and confocal microscopy.

## Author Contributions

Conceptualization: SB-R, CB-M, AIB

Methodology: SB-R, CB-M

Investigation: SB-R, CB-M

Visualization: SB-R

Supervision: CB-M, AIB

Writing—original draft: SB-R, CB-M

Writing—review & editing: SB-R, CB-M, AIB

