## Supplemental Figures for "A nematic framework for sprouting angiogenesis"

### 1 Supplementary Materials

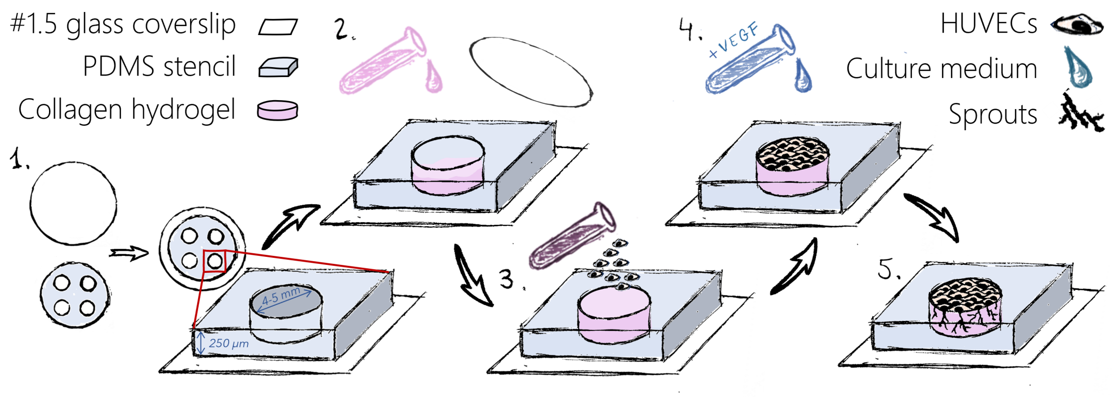


**Fig.** S1: **Protocol for the assembly of the flat system**. Steps: **1.** A PDMS stencil and a glass coverslip are plasma-bonded. **2.** A coverslip is placed on the hydrogel during polymerization to get a flat surface. **3.** Cell seeding is performed in a 12-well plate. **4.** The cell culture medium is supplemented with VEGF. **5.** Sprouts grow into the hydrogel.

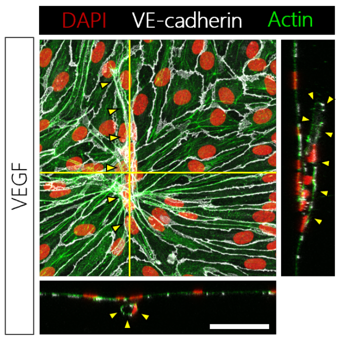


**Fig.** S2: **Example of lumenized sprout.** In-focus projection and cross sections (solid yellow lines denote the cross-sectional planes) of epifluorescence images of DAPI (red, nuclei), VE-cadherin (white, junctions) and phalloidin (green, actin) immunostaining reveal the existence of a lumen within a sprout. The sprout is denoted by yellow arrowheads. Scale bar, 50 µm.

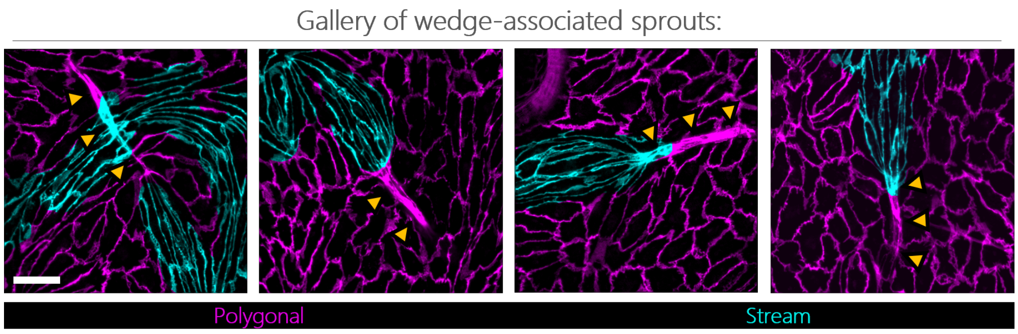


**Fig.** S3: **Examples of wedge-associated sprouts.** In-focus projection of epifluorescence images of VE-cadherin immunostaining. Sprouts are denoted by yellow arrowheads; cell stream and polygonal cell regions are shown in cyan and magenta, respectively. Scale bar, 50 µm.

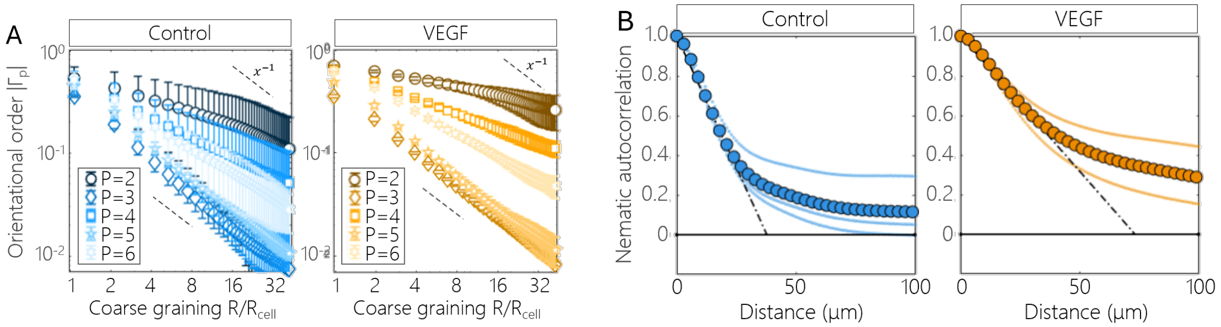


**Fig.** S4: **Control and VEGF conditioned monolayers exhibit nematic orientational order. A:** Magnitude of the coarse-grained p-atic orientational order $\Gamma_p$ as a function of the coarse-graining radius as calculated in Armengol-Collado et al. (71) (see Materials and methods). Data from n=3 independent experiments. Dashed lines represent a decay of $x^{-1}$, indicative of a purely random state. **B:** Nematic autocorrelation function calculated for control and VEGF-treated monolayers. The intersection of the tangent with the x-axis is considered to be the correlation length. VE-cadherin immunostaining images (n=3) were used to derive cell orientation.

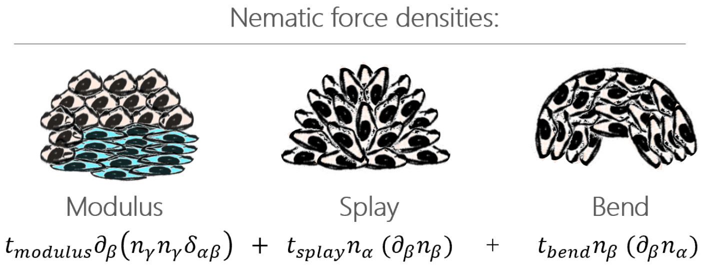


**Fig.** S5: **Illustrations of the modulus, splay and bend contributions to the traction force field in eq. 1.**

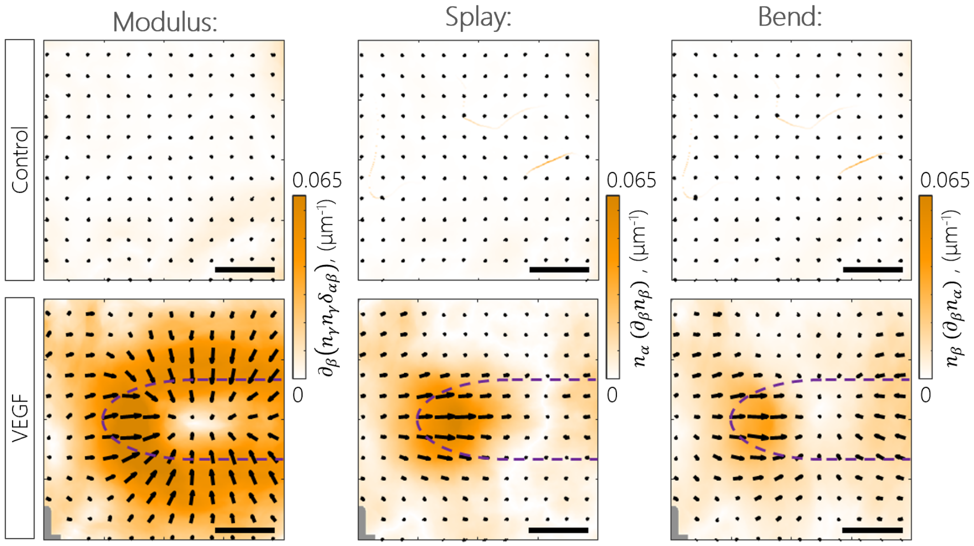


**Fig.** S6: **Modulus, splay and bend contributions to the nematic traction forces in the wedge region under VEGF conditioning.** Maps of the terms in eq. 1 proportional to $t_{modulus}$ (left panel), $t_{splay}$ (center panel) and $t_{bend}$ (right panel) corresponding to the data presented in fig. 4A (see Materials and methods). Pooled data ($n_{control}$=13 and $n_{VEGF}$=17) from $n_{control}$=4 and $n_{VEGF}$=7 independent experiments. Scale bars, 50 µm. Dashed lines indicate the approximate location of cell streams.

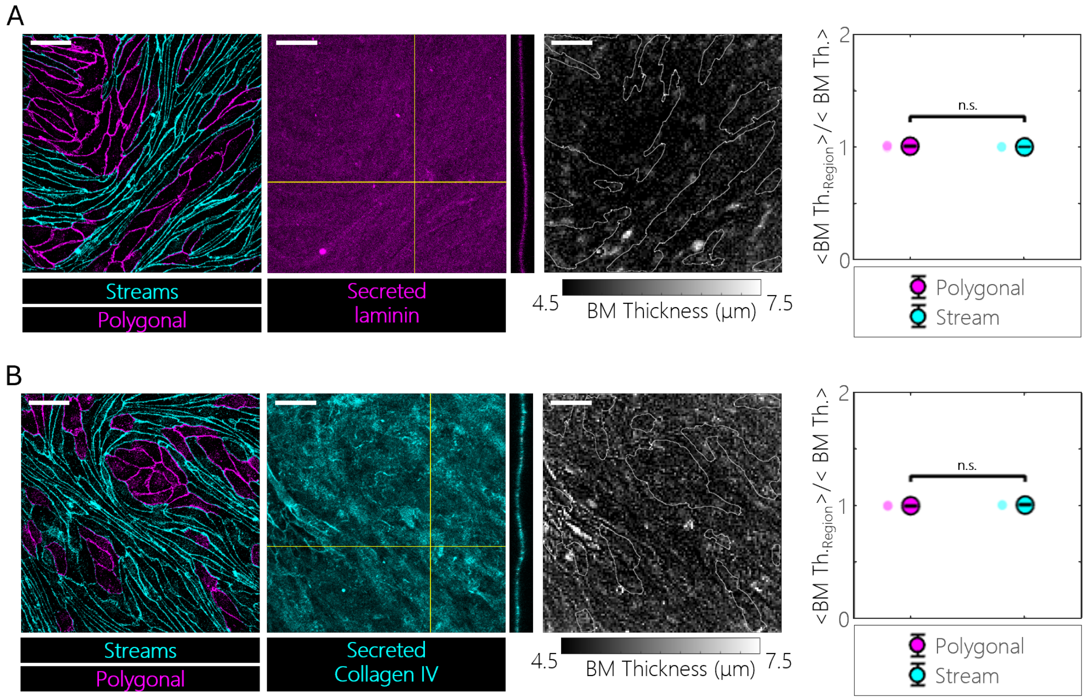


**Fig.** S7: **Deposition of laminin and collagen IV.** From left to right: Cell stream (cyan) and polygonal cell (magenta) regions based on VE-cadherin immunostaining of a VEGF-treated monolayer. Corresponding immunostaining for secreted basement membrane components-laminin (**A**, magenta), collagen IV (**B**, cyan) and cross-section (solid yellow lines denote the cross-sectional planes). Thickness map as obtained from confocal stacks, by calculating the full width at half maximum (FWHM) of a Gaussian fit of the Z-wise intensity for every XY coordinate. Quantification of the average regional thickness normalized by the total image average shows no significant difference among regions. Pooled data (n=3) from n=1 experiment per condition, two-sided Wilcoxon rank sum tests ($^{n.s.}P > 0.05$ represents statistical non-significance). Scale bars, 50 µm.

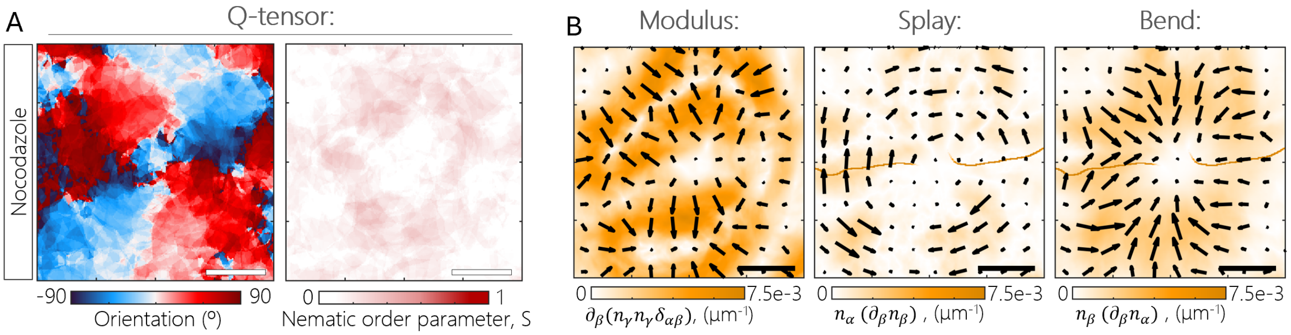


**Fig.** S8: **Contributions to the nematic traction forces under VEGF+nocodazole treatment. A:** From left to right, average orientation and nematic order parameter. **B:** Maps of the terms in eq. 1 proportional to $t_{modulus}$ (left panel), $t_{splay}$ (center panel) and $t_{bend}$ (right panel) corresponding to the data presented in panel **A** (see Materials and methods). Pooled data (n=20) from n=3 independent experiments. Scale bars, 50 µm.

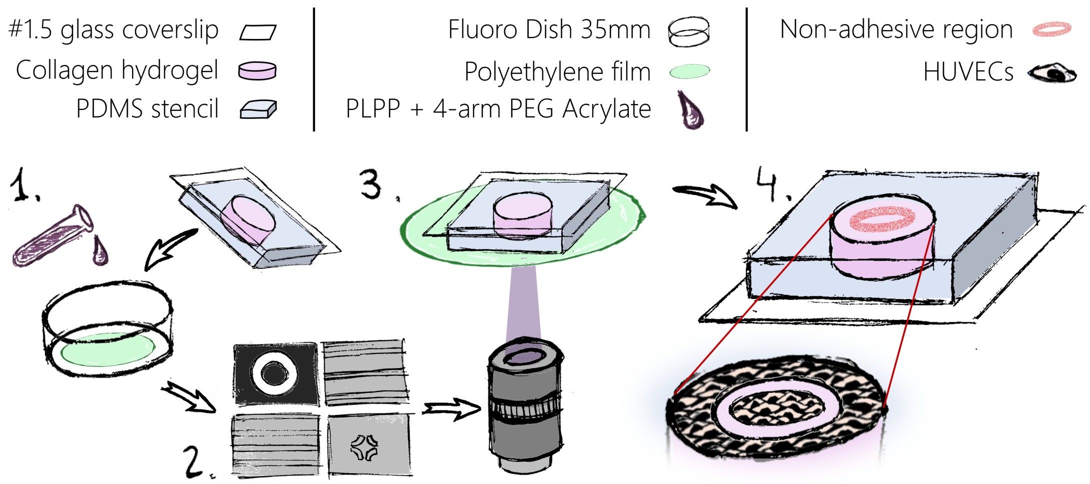


**Fig.** S9: **Protocol for the adhesive/non-adhesive patterning of collagen hydrogel surfaces.** Steps: **1.** A thin polyethylene (PE) film is placed inside a 35 mm-diameter Fluorodish, and a mixture of of PLPP photoinitiator and 4-arm PEG acrylate is deposited onto the PE film. **2.** The mixture is spread by placing the sample face-down, with repeated lifting and lowering to ensure penetration. **3.** UV illumination is applied at the PE-hydrogel interface following the desired shape of the non-adhesive region. **4.** HUVECs are seeded on the system, attaching only to the adhesive region.

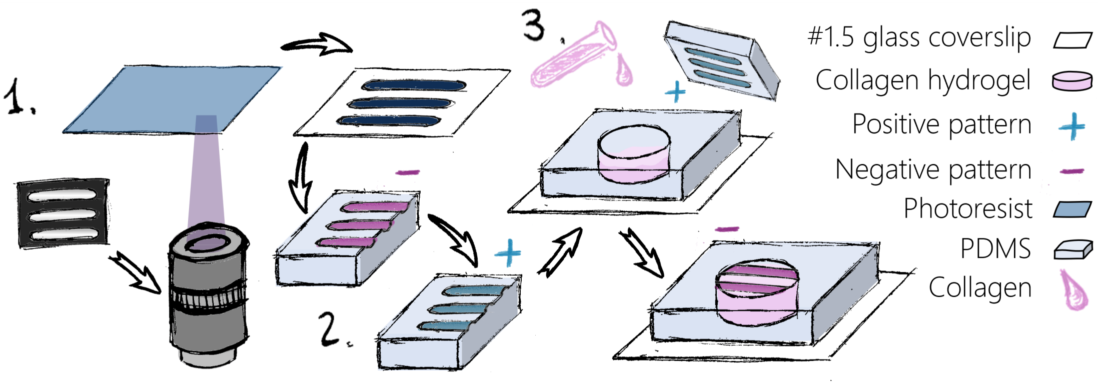


**Fig.** S10: **Protocol for the topographical patterning of collagen hydrogel surfaces.** Steps: **1.** Polymerization of a dry film photoresist following the desired pattern. **2.** Two PDMS casting steps to fabricate the final PDMS template. **3.** The PDMS template was placed on the hydrogel during polymerization.

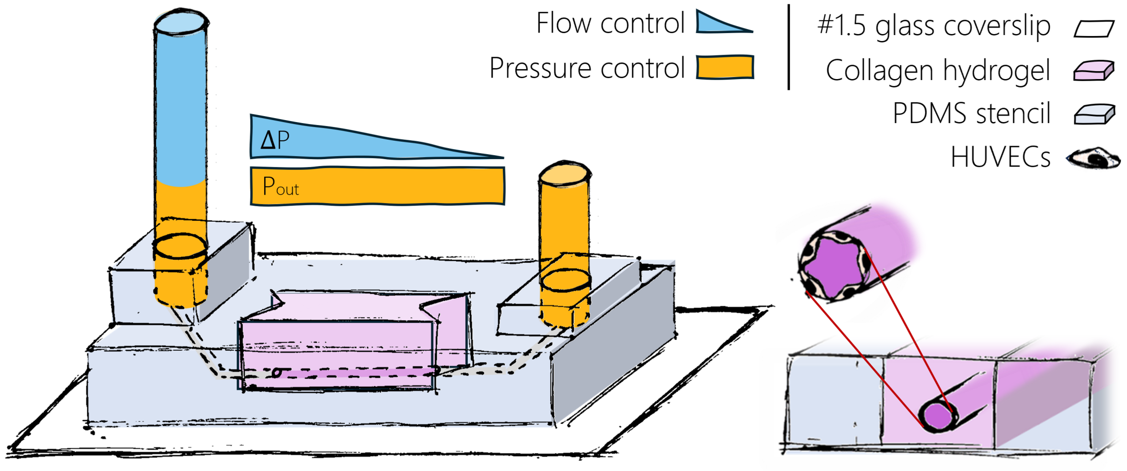


**Fig.** S11: **Microvessel-on-chip system schematics and working principle.** The system consists of a collagen hydrogel framed by PDMS. A 350 µm-diameter EC-lined channel is connected to two reservoirs. A syringe pump is connected to the inlet reservoir feeding it continuously. The height of the outlet reservoir imposes the pressure in the channel, while that of the inlet reservoir ensures the imposed flow rate.

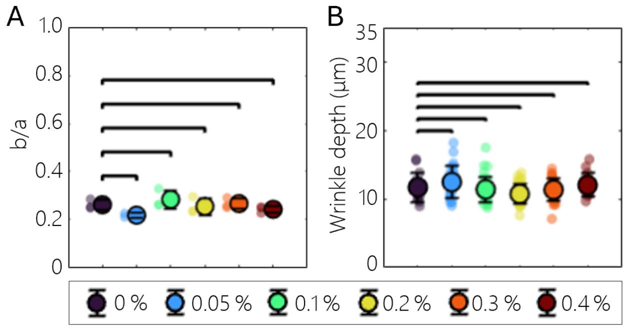


**Fig.** S12: **Effect of DMSO on cell shape and wrinkle depth. A:** Cell inverse aspect ratio and **B:** wrinkle depth in experiments containing VEGF and different doses of DMSO (n=3 independent experiments). Inverse aspect ratio data are averages from n=3 independent experiments, groups are compared using one-way ANOVA, Dunnett's post-test. Wrinkle depth data are pooled from n=3-7 independent experiments, groups are compared using Kruskal-Wallis, Dunn's post-test ($^{n.s.}P < 0.05$ represents statistical non-significance). Error bars represent ±SD.

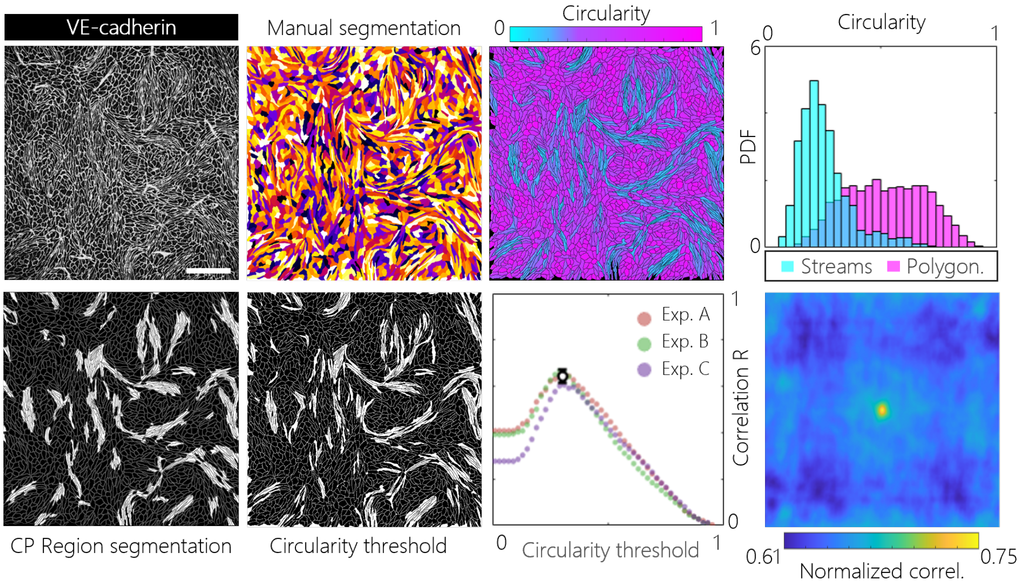


**Fig.** S13: **Comparative study of cell stream detection approaches.** The regional segmentation described in Materials and methods (1st column) is compared to an approach where manually segmented cells are attributed to cell stream or polygonal cell regions based on a circularity threshold (2nd column, and 3rd column, top). The threshold is selected by maximizing image correlation between the two segmentation methods (3rd column, bottom) and is similar for n=3 independent experiments. The distributions of cell circularity in the cell stream and polygonal cell regions detected with the selected methods show some overlap, indicating contributions of other factors to the Cellpose model, but show a clear trend (4th column, top). Segmentation from both strategies exhibits a high degree of spatial correlation (4th column, bottom), highlighting their compatibility in detecting cell streams and wedges. Scale bar, 250 µm.

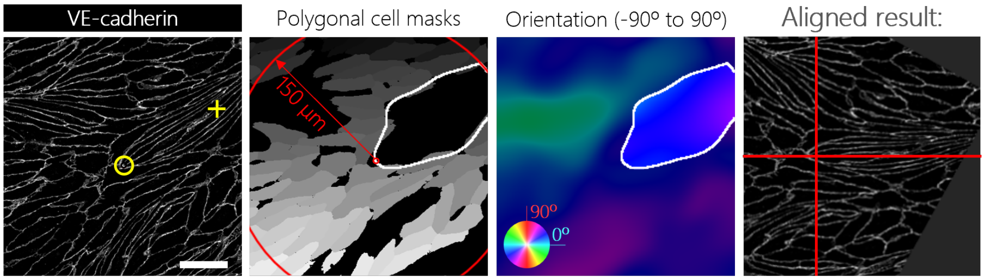


**Fig.** S14: **Semi-automatic alignment of images based on wedge position and cell stream direction.** From left to right: approximate wedge location (yellow circle) and stream direction (yellow cross) are manually provided. Cell streams are segmented and wedges are located as explained in Materials and methods. Stream orientation is calculated as the average orientation of cells inside the stream (only pixels within a distance of 150 µm to the wedge are considered). Images are displaced and rotated providing the resulting aligned VE-cadherin image (the red cross is centered on the wedge, and the cell stream is aligned with the horizontal axis). Scale bar, 50 µm.
